# L-Glyceraldehyde inhibits neuroblastoma cell growth via a multi-modal mechanism on metabolism and signaling

**DOI:** 10.1101/2023.12.20.572547

**Authors:** Martin Forbes, Richard Kempa, Guido Mastrobuoni, Liam Rayman, Matthias Pietzke, Safak Bayram, Birte Arlt, Annika Spruessel, Hedwig Deubzer, Stefan Kempa

## Abstract

Glyceraldehyde (GA) is a 3-carbon monosaccharide that can be present in cells as a by-product of fructose metabolism. Bruno Mendel and Otto Warburg showed that the application of GA to cancer cells inhibits glycolysis and their growth. This phenomenon was extensively studied up until the 1970’s. However, the molecular mechanism by which this occurred was not clarified. We describe a novel multi-modal mechanism by which the L-isomer of GA (L-GA) inhibits cancer cell growth. L-GA induces significant changes in the metabolic profile, promotes oxidative stress and hinders nucleotide biosynthesis. GC-MS and ^13^C-labelling was employed to measure the flow of carbon through glycolytic intermediates under L-GA treatment. It was found that L-GA is a potent inhibitor of glycolysis due to its proposed targeting of NAD(H)-dependent reactions. This results in growth inhibition, apoptosis and a redox crisis in the cancer cell. It was confirmed that the redox mechanisms were modulated via L-GA by proteomic analysis. This elucidated a specific subset of proteins harbouring oxidoreductase and antioxidant activity. Analysis of nucleotide pools in L-GA treated cells depicted a remarkable and previously unreported phenotype. Nucleotide biosynthesis in neuroblastoma cells is significantly inhibited upon L-GA treatment. Through the application of the antioxidant N-acetyl-cysteine in conjunction with L-GA, metabolic inhibition was partially relieved. We present novel evidence for the multi-modal mechanism of L-GA action in neuroblastoma cells. Specifically, a simple sugar that inhibits the growth of cancer via dysregulating the fragile homeostatic environment inherent to the cancerous cell.

## Introduction

In 1929 Bruno Mendel reported that glyceraldehyde (GA) inhibits anaerobic fermentation in Jensen sarcoma without affecting respiration in tumor or normal rat tissue (1). Glyceraldehyde is the non-phosphorylated form of the glycolysis intermediate, glyceraldehyde-3-phosphate. Five years previous to Mendel’s discovery Otto Warburg found that cancer cells exhibit an addiction to glucose and show high glycolytic activity. Specifically, cancer cells have a preference towards glycolysis and secretion of lactate over oxidative phosphorylation even in the presence of oxygen (2). This observation between the metabolism of non-cancerous and cancerous cells gives reasoning to the efficacy of GA as a therapeutic, via glycolytic inhibition.

This prompted the seminal research led by Warburg, showing that GA was effective in curing ascites cancer in all 75 mice that were treated (3). Numerous laboratories conducted experiments with GA and reported varied efficacy, or none at all (4, 5, 6). A driving factor in the contentious nature of GA efficacy is that, the D-and L-isomer of GA had differing potency (7). The synthesis of pure L-GA demonstrated its superiority over D-GA in glycolytic inhibition (8).

Due to only relative success *in vivo*, GA went largely out of fashion. One of the last published experiments into the efficacy of GA *in vivo* showed that neuroblastoma cells were sensitive to GA and exhibited inhibited glycolysis and cell proliferation (9). However, the molecular mechanism by which GA inhibits cancer cells was not established. Later research focused solely on GA as a stimulant of insulin secretion in pancreatic cells, thereby shelving its potential as a therapeutic (10, 11).

Rapid but inefficient production of ATP via glycolysis comes at the cost of an increased oxidative state (12). Cancer cells in proliferative states require a metabolic adaptation to accommodate the in-crease of reductive and oxidative cellular processes (redox). Reactive oxygen species (ROS) accumulate due to the increase in metabolic processes. The cancer cell requires mechanisms to detoxify the cell of ROS and maintain a homeostatic redox balance (13). This system is finely tuned to avoid ROS induced autophagy and apoptosis (14). By targeting ROS sensitive metabolic processes including: glycolysis and nucleotide biosynthesis, opportunities for novel - or a renaissance of - therapeutics arise.

Neuroblastoma is an extra-cranial solid tumour occurring in the developmental sympathetic nervous system. Of all childhood cancers, neuroblastoma accounts for 15%, making it the most commonly diagnosed malignancy in the first years of life (15). Approximately 90% of neuroblastoma tumours are diagnosed in children under the age of 10, with a median age of 18 months (16). In high risk groups long-term survival is below 50%. In addition, the treatment regimens are severe and invasive with chemotherapy, surgery, radiation and stem cell transplantation being common (17, 18).

A common feature of *MYCN* amplified neuroblastoma cells is their high levels of ROS production (19). Given that MYCN increases ROS production, compounds that further increase ROS may provide valid targets for therapeutics. As a pretext to the research presented here, it was found that GA had the +potential for generating reactive oxygen species in pancreatic islet cells (20). There is scope to assess the function of L-GA in relation to ROS production and its efficacy in tipping the fine balance towards cancer cell death.

We propose that L-GA acts in a multi-modal fashion on metabolism to inhibit cancer cell growth. Here we examine effect of L-GA on metabolic and proteomic processes in neuroblastoma and fibroblast cells. The data alludes to the generation of ROS, in combination with inhibitory action on central carbon metabolism and nucleotide biosynthesis pathways.

## Results

### L-Glyceraldehyde inhibits cell growth in neuroblastoma cells and induces apoptosis

The working concentration of glyceraldehyde was reported to be between 0.5 and 2 mM *in vitro* and 0.5- 1 g/kg *in vivo* (4, 5, 6, 9). We aimed to confirm the working concentration of L-GA in neuroblastoma cells and characterise the rate of growth inhibition. Five neuroblastoma cell lines were treated with a range of L-GA concentrations (0-10 mM) for 24-96 hrs and a WST-1 assay was performed. The IC50 of L-GA was found to be in the range of 262-1005 µM after 24 hrs treatment (supplementary Table 1)). Additionally, neuroblastoma cell lines: IMR-32 and SH-SY5Y and a non-cancer fibroblast cell line: VH-7, were treated with a range of L-GA concentrations (0-1.5 mM) for 24 hrs prior to cell counting. VH-7 cells did not achieve an 50% decrease at the highest L-GA concentration used in this experiment. IMR-32 and SH-SY5Y both achieved a similar 50% decrease in cell number at L-GA concentrations: 0.76 mM and 0.55 mM respectively (Figure 1A). These results are similar to the L-GA concentrations found to inhibit glycolysis during experiments conducted in the 1940s, whereby sub 1 mM L-GA inhibited glycolysis in muscle extracts (4).

**Figure 1.**
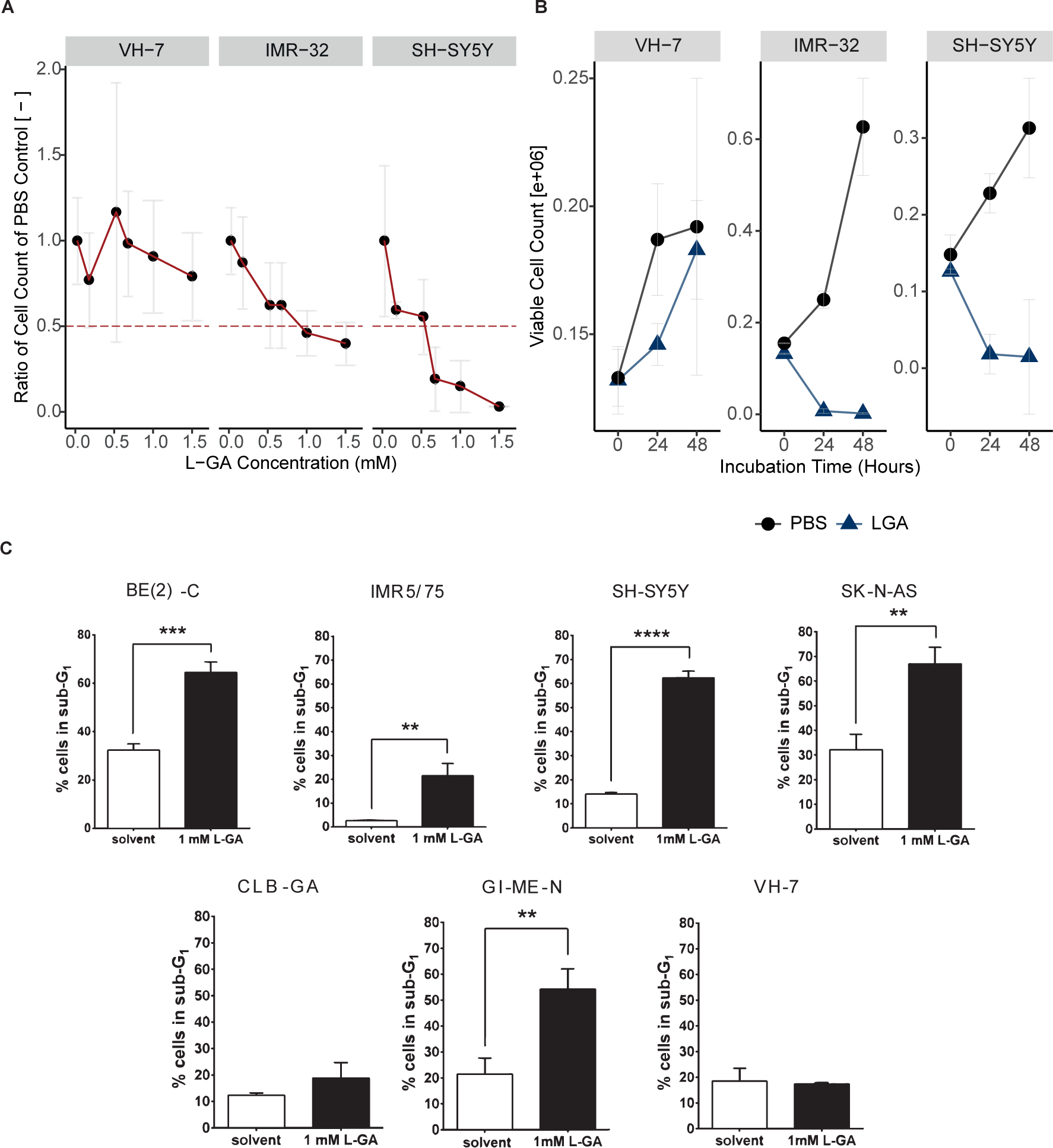
L-Glyceraldehyde inhibits cell growth in neuroblastoma cells. A Cell lines VH-7, IMR-32 and SH-SY5Y cells were incubated with varying concentrations of L-GA for 24 hours after seeding. Viable cell counts were taken for each L-GA concentration via the trypan exclusion method and normalised to 0.0 mM control. The 50% reduction in cell count is denoted by a dotted red line with errors bar representing the ± SEM (n=3). B Cell lines were incubated with 1 mM L-GA (blue) or PBS (black) for 48 hrs after seeding and settling for 24 hrs. Viable cell counts were taken every 24 hrs via the trypan exclusion method. Error bar representing the ± SE of the mean (n=3). C Cell lines BE(2)-C, IMR5/75, SH-SY5Y, SK-N-AS, CLB-GA, GI-M-EN and VH-7 were treated with L-GA for 72 hrs before flow cytometry analysis. Sub-G1 cells were gated and a two-sided T-test was performed (n=3).

In order to assess the effect of L-GA on cell proliferation, 1 mM L-GA was applied to VH-7, IMR-32 and SH-SY5Y for up to 48 hrs (Figure 1B). Viable cell counts were measured every 24 hrs. Following 24 hrs of incubation with L-GA, cell counts were reduced approximately 10-fold relative to the control in IMR-32 and SH-SY5Y. Inhibition of cell growth persisted over the 48 hrs incubation. The fibroblast cell line, VH-7, showed a less extreme response to L-GA treatment than the neuroblastoma cell lines, however the cells grow slower in the presence of L-GA.

Flow cytometry was performed to examine whether apoptosis is induced by L-GA. The sub-G1 fraction of a panel of L-GA treated cells was measured following 72 hrs exposure (Figure 1C). All neuroblastoma cell lines except CLB-GA presented a significant increase in the sub-G1 phase, alluding to an apoptotic response. VH-7 did not show an increase in the sub-G1 fraction in response to L-GA.

We concluded that L-GA is active against neuroblastoma cells at concentrations *≤* 1 mM. Sub-G1 cell cycle fractions were indicative of apoptotic response and motivated us to perform further multi-omic experiments to identify the mode of L-GA action on cell death.

### Proteomic analysis of L-GA treated NB cells reveals an increase of proteins associated with oxidoreductase activity

At present, no literature reports the effect of L-GA on the proteome. In order to garner a broad picture of the effect of L-GA, LC-MS shotgun proteomics was performed in five neuroblastoma cell lines (SH-SY5Y, IMR-32, BE(2)-C, GI-M-EN, SK-N-AS) and one non-cancerous fibroblast cell line (VH-7). Cells were challenged with 1 mM L-GA for 24 hrs. Proteins were extracted from cell lysates and label free quantification was performed. Label free quantities (LFQ) of 3,551 proteins were measured, of which 244 were annotated to be associated with 6 pathways of interest. Hierarchical clustering (Figure 2A), of the difference in log2(LFQ) (L-GA-PBS) for each cell line was performed. Specifically, we aimed to investigate the proteins associated with: Apoptosis, Cell Cycle, Glycolysis, Nucleotide Metabolism, Oxidoreductase and the TCA cycle. SH-SY5Y and IMR-32 showed the strongest increase in proteins of the oxidoreductase family following L-GA treatment. GI-M-EN and SK-N-AS also showed increased oxidoreductase activity whereas VH-7 and BE(2)-C produced a less pronounced response in the same pathway.

**Figure 2.**
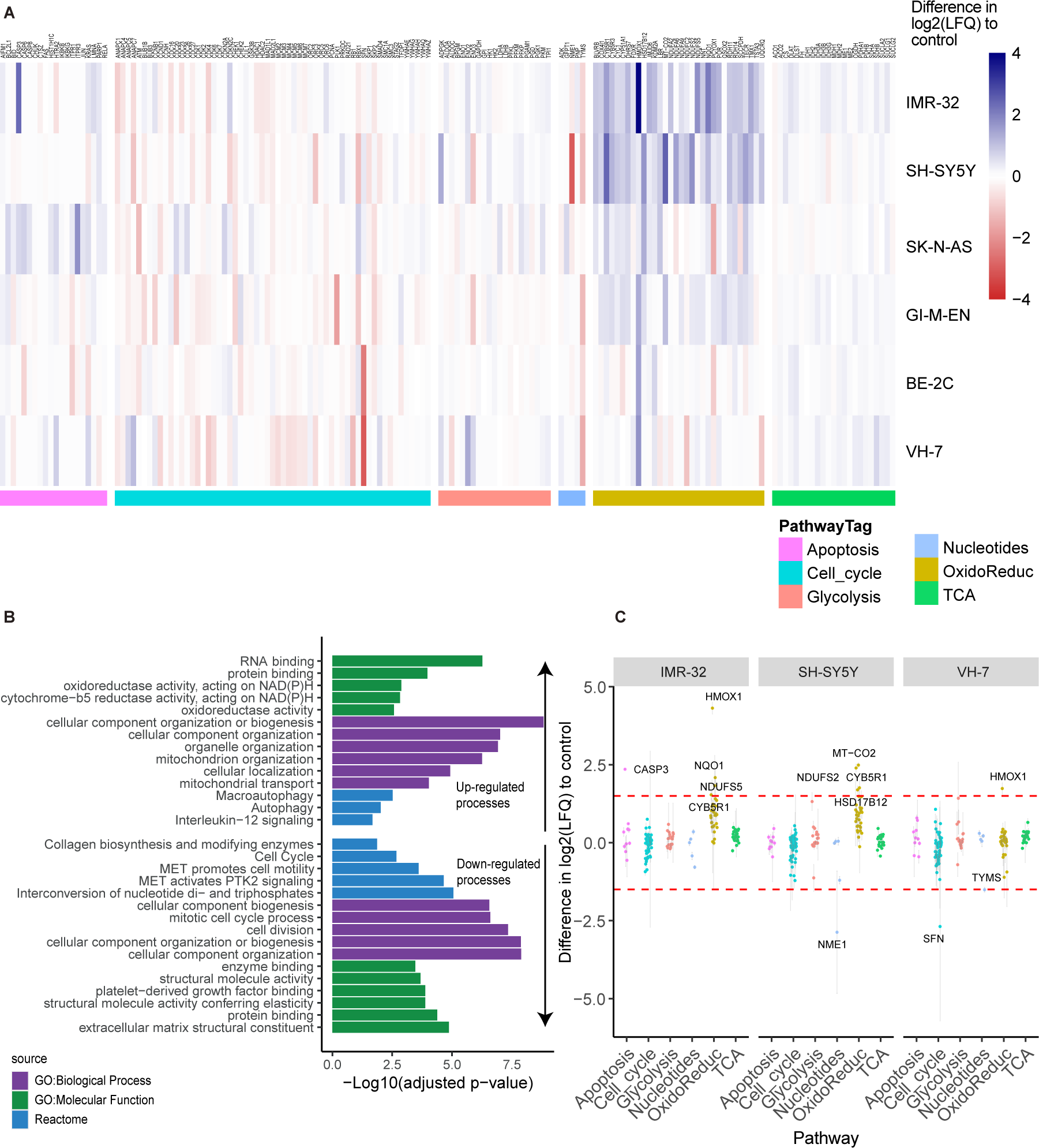
Proteomic analysis of L-GA treated NB cells. A Proteomic analysis of cell lines: BE-2C, GI-M-EN, VH-7, SH-SY5Y, IMR-32 and SK-N-AS treated for 24 hrs with 1 mM L-GA or PBS and subjected to LC-MS proteomic analysis. Hierarchical clustering of cell lines show clustering of pathways between cell types. Data represents the difference in Log_2_ label free quantity Log_2_(LFQ) of L-GA treated cells relative to PBS treated cells, n=3 for each cell line. **B** GO enrichment analysis of data filtered by > 1 and < -1 difference Log_2_(LFQ) of L-GA treated cells relative to PBS control in all cell lines. The top 5 term names, defined by lower adjusted p-value < 0.05, of GO:Molecular function, GO:Biological process and REACTOME are presented. **C** Proteins from figure A were filtered by > 1.5 and < -1.5 difference Log_2_(LFQ) of L-GA treated cells relative to PBS in IMR-32, SH-SY5Y and VH-7. Filtered proteins are highlighted by a red dotted line. Proteins that show differences outside of this line are labelled by text.

Ontological enrichment was performed on all proteins in which the difference in log2(LFQ) (L-GA-PBS) was > 1 or < -1 in order to define which molecular and biological processes were perturbed upon L-GA treatment in all cell lines (Figure 2B) (21). The top 5 term names, defined by lower adjusted p-value *≤* 0.05, of: GO:Molecular function, GO:Biological process and REACTOME were selected for graphical representation. Up-regulated, defined by a difference of > 1, GO subsets showed enrichment of oxidoreductase activity, cellular organisation and autophagy. Down-regulated process, defined as a difference < -1, were found to be enriched in the: cell cycle and cell division processes, inter-conversion of nucleotide di and tri-phosphates and extracellular matrix constituents. We selected SH-SY5Y and IMR-32 from the hierarchical clustering data to compare against the fibroblast cell line, VH-7. It was found that, in general, proteins of the oxidoreductase family showed the greatest increase in IMR-32 and SH-SY5Y when exposed to L-GA. HMOX1 was found to be 4-times higher in IMR-32 (Figure 2C).

We concluded that L-GA causes modulation to oxidative stress response, nucleotide biosynthesis and cell cycle proteins. Further experiments aimed to characterise the metabolic mode of action on these processes.

### L-GA depletes nucleotides pools, causes cytoskeletal aberrations, cell cycle arrest and inhibition of growth

Glycolytic inhibition via L-GA has been extensively reported and challenged since the 1930’s (2, 4, 5, 3, 22). It was hypothesised that L-GA would deplete ATP and total nucleotide pools. The depletion of nucleotide pools results in a stalling of DNA synthesis and cell cycle arrest. Furthermore, the imbalance of purine and pyrimidine nucleotide pools induce pro-apoptotic pathways and inhibit cell proliferation (23, 24). NAD(P)+ and NAD(P)H availability is essential to maintaining the redox state of the cell and adequate performance of NAD-dependent enzymes.

The *de novo* synthesis of nucleotides originates from the production of ribose-5-phosphate (R5P) derived from the oxidative or non-oxidative pentose phosphate pathway (PPP). The non-oxidative pathway produces R5P through the cycling of carbons from F6P and GA3P. It is of particular interest that the non-oxidative pathway requires GA3P as a substrate. We suspected that L-GA perturbs the synthesis of nucleotides via disruption of R5P synthesis, thereby inducing limited nucleotide availability for correct cell cycle maintenance.

Initial experiments followed previous protocols, whereby IMR-32, SH-SY5Y and VH-7 were treated with 1 mM L-GA for 24 hrs. Cells were harvested, metabolites were extracted and enriched for nucleotides on a C18 stationary phase, and measured via DI-MS/MS. In neuroblastoma cells DNA and RNA nucleotide pools were significantly depleted following 24 hrs incubation with 1 mM L-GA (*p ≤* 0.0001) (Figure 3A). VH-7 showed less reduction in DNA and RNA nucleotide pools compared to the neuroblastoma cell lines.

**Figure 3.**
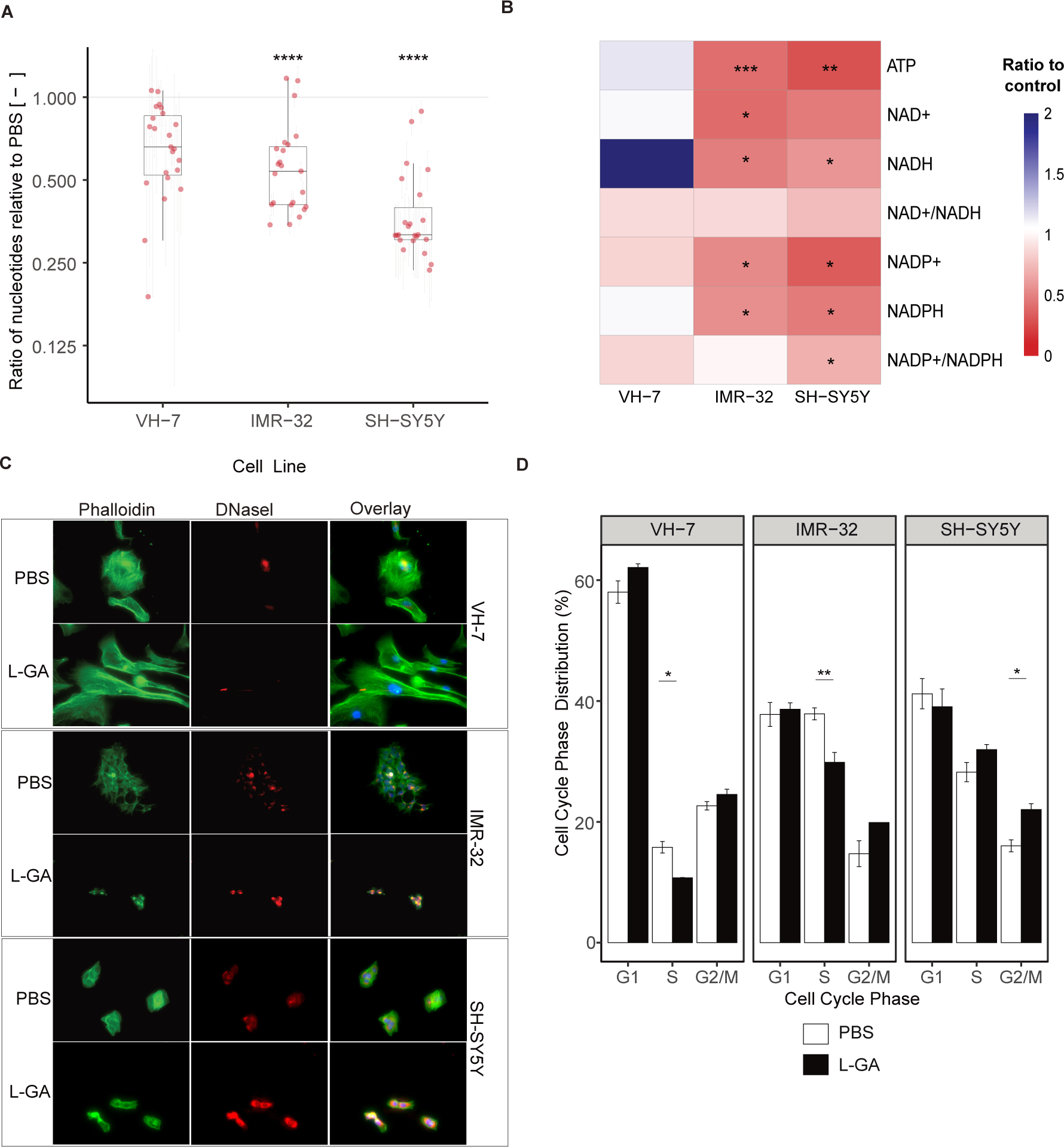
L-GA causes cytoskeletal aberrations and cell cycle arrest within 24 hrs. A Cell lines VH-7, IMR-32 and SH-SY5Y were treated with 1.0 mM LGA or PBS L-GA for 24 hrs. Nucleotides were extracted and measured by DI-MS/MS. Boxplots represent the mean ratio in each DNA and RNA nucleotide intensity normalised to PBS (n>3). Error bars represent ± SEM of each nucleotide measured. P-values were calculated using a two-tailed T-test. B Cells were treated as in A; Intensities of ATP, NAD+, NADH, NADP+ and NADH with associated ratios were calculated and normalised to PBS (n>3). P-values were calculated using a two-tailed T-test. **C** SH-SY5Y, IMR-32 and VH-7 were treated with 1 mM L-GA or PBS for 24 hrs and subjected to flow cytometry analysis. Cells were gated for cell cycle phases: G_1_, S and G_2_/M. The mean of the percentage of cells in each phase was calculated (n=3). Error bars represent *±* SEM. Two-tailed T-tests were conducted in each phase between treatments. **D** Cell lines VH-7, IMR-32 and SH-SY5Y were fixed to glass slides and stained with Phallodin-iFluor(F-) or Deoxyribonuclease I (G-) actin probes then imaged at 40X magnification using Keyence Bz-x700 microscope. Images are exemplary of 3 independent slides.

ATP, NAD+, NADH, NADP+ and NAPDH with associated ratios, of the cell were measured in VH-7, IMR-32 and SH-SY5Y (Figure 3B). In both neuroblastoma cell lines the ATP ratio significantly decreased (*p ≤* 0.01), this was not observed in VH-7. NAD(P)+ and NADP(H) were decreased in neuroblastoma cells with a slight decrease in the NAD+/NADH ratio. However it is to be noted that NAD+/NADH ratios were calculated when both co-factors are strongly depleted. NAD+ was depleted more in IMR-32 and NADP+ in SH-SY5Y.

L-GA caused an imbalance in the NAD+/NADH ratio at concentrations above 0.5 mM, in a similar fashion to total nucleotide pool depletion. The phosphorylation potential (ATP + 1/2 ADP / ATP + ADP + AMP) (25) of the cell is severely inhibited at 1 mM L-GA (supplementary Figures 9A,B). Taken together, it is apparent that the cell experiences redox stress and a reduction in the ability to perform phosphorylation reactions.

We suspected that the changes in cell morphology shown by phase-contrast microscopy (not shown), is a result of the dysregulation in the maintenance of the ATP-dependent microfilaments of the cytoskeleton. A mature cytoskeletal microfilament is composed of sub-units of actin (G-actin) polymerised into actin filaments (F-actin). The maintenance of the F-actin polymer is an ATP-dependent reaction. Therefore, the inhibitory action of L-GA on glycolysis and ATP production may perturb actin polymerisation.

In order to assess the effect of L-GA on the cytoskeleton, cells were stained simultaneously for F-actin and G-actin. Briefly, cell lines: IMR-32, SH-SY5Y and VH-7 were treated with 1 mM L-GA for 24 hrs. Slides were imaged via fluorescent excitation and emission (ex/em) 493/517 nm for phalloidin (F-actin) and 590/617 nm for DNase I (G-actin) (Figure 3C, supplementary Figures 9H). Image analysis showed increased G-actin fluorescence and a dysregulation of F-actin structures in neuroblastoma cells. VH-7 fibroblast cell line, actin structure appeared to be less compromised upon L-GA treatment. Given that cellular division is a process in which regulation of the cytoskeleton is essential, it was assessed whether cellular division is also compromised.

Cells were treated with 1 mM L-GA for 24 hrs before harvesting and stained with propidium iodide and analysed by flow cytometry. Cells were gated into G1, S or G2/M cell cycle phases and the distribution was calculated in percentage. Cell lines: VH-7, IMR-32 and SH-SY5Y investigated as shown in Figure 3D. IMR-32 showed a significant decrease in cells in the S phase (*p ≤* 0.01) and an increase in the G2/M phase. SH-SY5Y cells observed an increase in the S-phase and an increase in the G2/M phase (*p ≤* 0.05). In VH-7 there was a minute yet increase in the G2/M phase and a significant decrease in the S phase in response to L-GA (*p ≤* 0.05).

The cell cycle arrests that neuroblastoma cells experience lead to the hypothesis that essential nutrients aren’t available to the cell. The regulation of the cell cycle is tightly controlled. Checkpoint kinases receive input from nutrient levels to initiate entry and exit from cell cycle phases (26, 27, 28, 29). Consequently, we aimed to assess the effect of L-GA on glycolysis and nucleotide metabolism.

### L-GA rapidly inhibits glycolysis and nucleotide metabolism

Previous research showed that L-GA inhibited glycolysis of T98G and HEK293 cells (30). Specifically, L-GA treatment resulted in the inhibition of the flux of carbon from glucose into metabolites downstream of aldolase-which utilises glyceraldehyde-3-phosphate (GA3P) as a substrate. Furthermore, L-GA was shown to be more effective at inhibiting glycolysis than D-GA chiral isomer. Given this, the aim was to show that neuroblastoma cells are also susceptible to glycolytic inhibition via L-GA. To measure the flux of carbon through metabolic pathways we measured the change in metabolite pool sizes in cooperation with pulsed stable isotope resolved metabolomics (pSIRM) to derive the quantity of ^13^C-labelled metabolites (31). We aimed to characterise the effect of L-GA in the early stages of treatment time on central carbon metabolism and nucleotide metabolism.

Initially we assessed how rapidly nucleotides and NAD(P)/H species deplete following L-GA treatment. IMR-32 and SH-SY5Y were treated for 1 hr and 8-hrs and nucleotides were extracted and measured as previously described (Figure 4A). SH-SY5Y cells appeared to increase ATP, NAD(H) and NADP(H) after one hour - with NAD+ showing a strong increase - yet early nucleotide intermediates start to decrease (R5P, PRPP). In IMR-32 nearly all nucleotides are slightly decreased after 1 hr treatment, except for AICAR and NADPH. Following 8 hours treatment, almost all nucleotides are severely decreased in both cell lines. In addition we found that D-GA was less effective at depleting nucleotides than L-GA even after 24 hrs in both cell lines (supplementary Figures 9D). Given this, we conducted pSIRM after 1 hr treatment to capture early metabolic effect.

**Figure 4.**
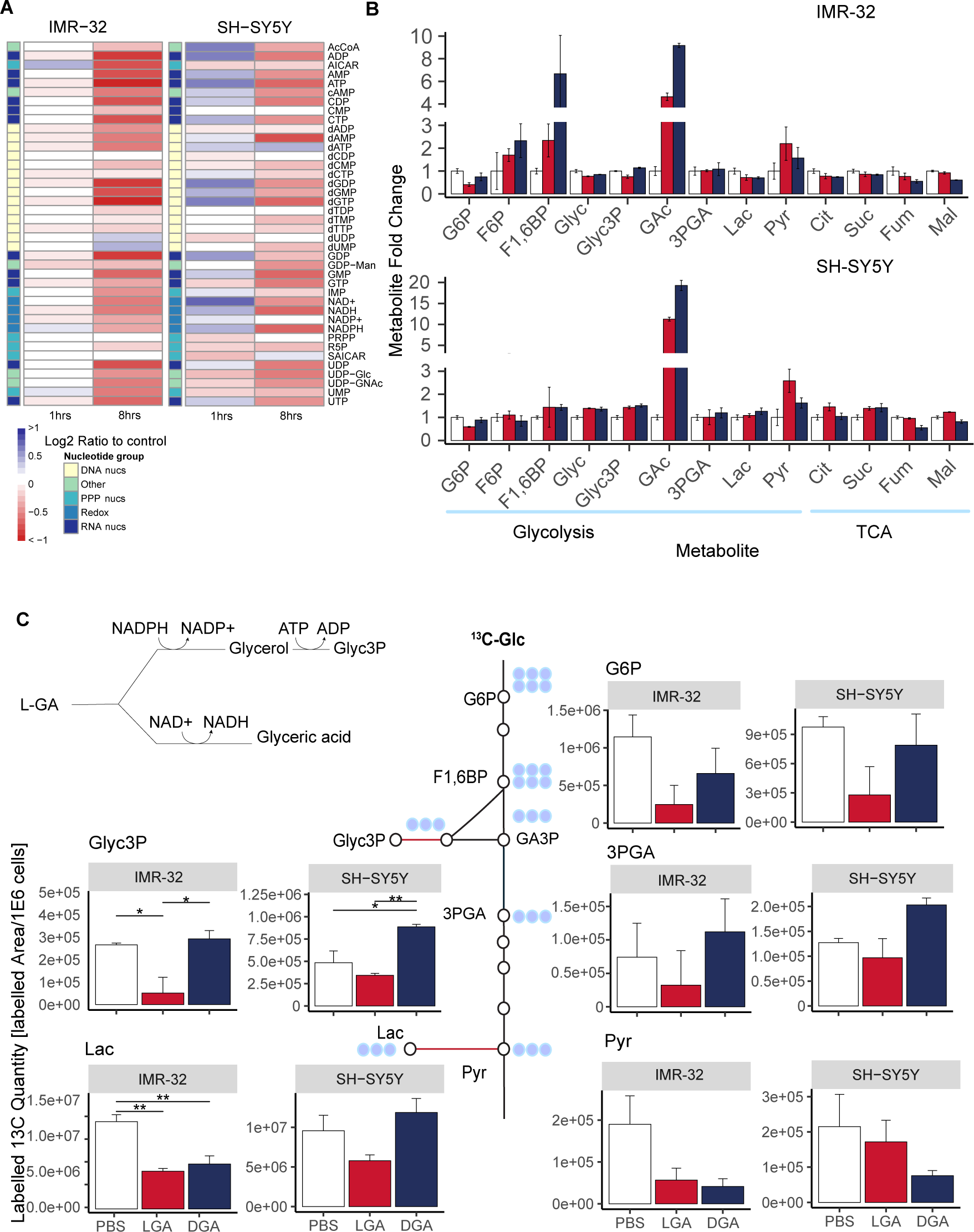
L-GA selectively and rapidly inhibits glycolysis. A Cell lines IMR-32 and SH-SY5Y were treated with 1 mM L-GA for 1 or 8hrs. Nucleotides were extracted and measured by DI-MS/MS. The fold change in nucleotide intensity relative to PBS was calculated for representation by heatmap (n=4). B Polar metabolites were extracted from IMR-32 and SH-SY5Y cell lines following treatment of 1 mM L-GA or 1 mM D-GA for 1 hr. Samples were processed via GC-MS and analysed to attain the abundance of metabolites relative to PBS. Bar charts represent the metabolite mean fold change, error bars represent ± SEM (n=3). C Cell lines IMR-32 and SH-SY5Y were treated with 1 mM L-GA or 1 mM D-GA for 1 hr and labelled for 10 mins with U-13C-glucose within the treatment time. Bar charts depict the mean labelled 13C-quantity (Labelled peak area/1E6 cells) of depicted metabolites. Error bars represent ± SEM (n=3). P-values were calculated using Anova and Tukey post-hoc test.

Cell lines IMR-32 and SH-SY5Y were treated for 1 hour with 1 mM L-GA or 1 mM D-GA and labelled with ^13^C-glucose for 10 mins within the treatment time. Extracted metabolites were analysed by GC-MS for their quantity and amount of ^13^C-label. In both cell lines the quantity of glyceric acid (GAc) was significantly increased upon both L-GA and D-GA treatment (Figure 4B). In IMR-32 Glycerol (Glyc) did not show the same increase, suggesting that glyceraldehyde is converted to, mostly, GAc. This process consumes NAD(P)+ via aldehyde dehydrogenase. However, in SH-SY5Y more Glyc was produced than in IMR-32. This process occurs via aldehyde reductase, consuming NAD(P)H. In IMR-32 and SH-SY5Y there was no increase in glyceric acid-3-phosphate (3PGA) suggesting GAc cannot be phosphorylated further. In SH-SY5Y, there is an increase in glycerol-3-phosphate (Glyc3P), converse to IMR-32. The first step in glycolysis, glucose (Glc) to Glucose-6-phosphate (G6P) shows reduced G6P quantities in both cell lines, only when L-GA is present. Similarly, L-GA induces greater pyruvate (Pyr) accumulation than D-GA. In IMR-32 TCA cylce intermediates are slightly decreased in both L-GA and D-GA conditions. In SH-SY5Y citrate (Cit), succinate (Suc) and malate (Mal) are slightly increased and fumarate (Fum) is slightly decreased under L-GA treatment.

Figure 4C shows ^13^C-labelled quantities (^13^C-Peak area/1E6 cells) of glycolytic intermediates following 10 mins labelling with ^13^C-glucose. In both cell lines G6P is depleted, however only with L-GA treatment. Labelled quantities of 3PGA is decreased in both cell lines upon L-GA treatment, although not with D-GA. Label incorporation into Glyc3P is significantly depleted in IMR-32 although only with L-GA, SH-SY5Y does not show the same degree of depletion. Importantly, D-GA does not deplete Glyc3P labelled quantities in either cell line. Labelled quantities of Lac are reduced significantly by L-GA and D-GA, although only in IMR-32. Pyr shows decreased labelled ^13^C quantities, however this is more apparent in IMR-32.

Given that glycolysis was decreased we examined as to whether the cell would fuel the TCA via glutaminolysis. Cell lines IMR-32 and SH-SY5Y were treated for 1 hour with 1 mM L-GA or 1 mM D-GA and labelled with ^13^C-glutamine for 30 mins within the treatment time, supplementary Figures 9E), in both cell lines we found highly increased labelled ^13^C-quantities in TCA intermediates when applying L-GA. In IMR-32, D-GA did not produce the same response as L-GA, however in SH-SY5Y D-GA also increased labelled TCA intermediates. The increase in mitochondrial activity increases reactive oxygen species (ROS). We suspected that the L-GA induced glutaminolysis promotes oxidative stress via ROS generation.

### L-GA induces ROS production, inhibiting nucleotide synthesis and arresting cell cycle progression

We aimed to quantify the generation of reactive oxygen species in L-GA treated cells. This was prompted by the observation of dysregulated NAD species, and the increase in the response of the oxidoreductase pathway in the proteomics data. We suspected that the co-application of an antioxidant, N-acetyl-cysteine (NAC) would relieve ROS dependent cell stress.

Cell growth analysis was performed over a 72 hr time period with treatments of: 1X PBS, 1 mM L-GA, 100 µM H_2_O_2_, 1 mM L-GA + 5 mM NAC, 100 µM H_2_O_2_ + 5 mM NAC and PBS + 5 mM NAC. Figure 5A shows cell counts sampled every 24 hrs in duplicates for IMR-32 and SH-SY5Y. For both cell lines, cell growth is inhibited with 1 mM L-GA. Upon supplementation with 5 mM NAC, L-GA treated cells showed less severe and delayed growth inhibition. 100 µM H_2_O_2_ stopped cell growth in IMR-32 cells within 24hrs to below detectable cell concentrations. The addition of NAC recovered the effect of H_2_O_2_ on cell growth. SH-SY5Y showed markedly less inhibition of cell growth in response to H_2_O_2_ compared to IMR-32. In both cell lines, NAC only partially recovered cell growth in L-GA treated cells whereas NAC negated the effect of H_2_O_2_, indicating that oxidative stress induced by L-GA is only one facet of its action. Furthermore, applying L-GA to cells and replacing with fresh media did not result in cellular recovery (supplementary Figures 9C). To clarify this, we aimed to show that the inhibitory effect of L-GA was in partial relation to the production of ROS.

**Figure 5.**
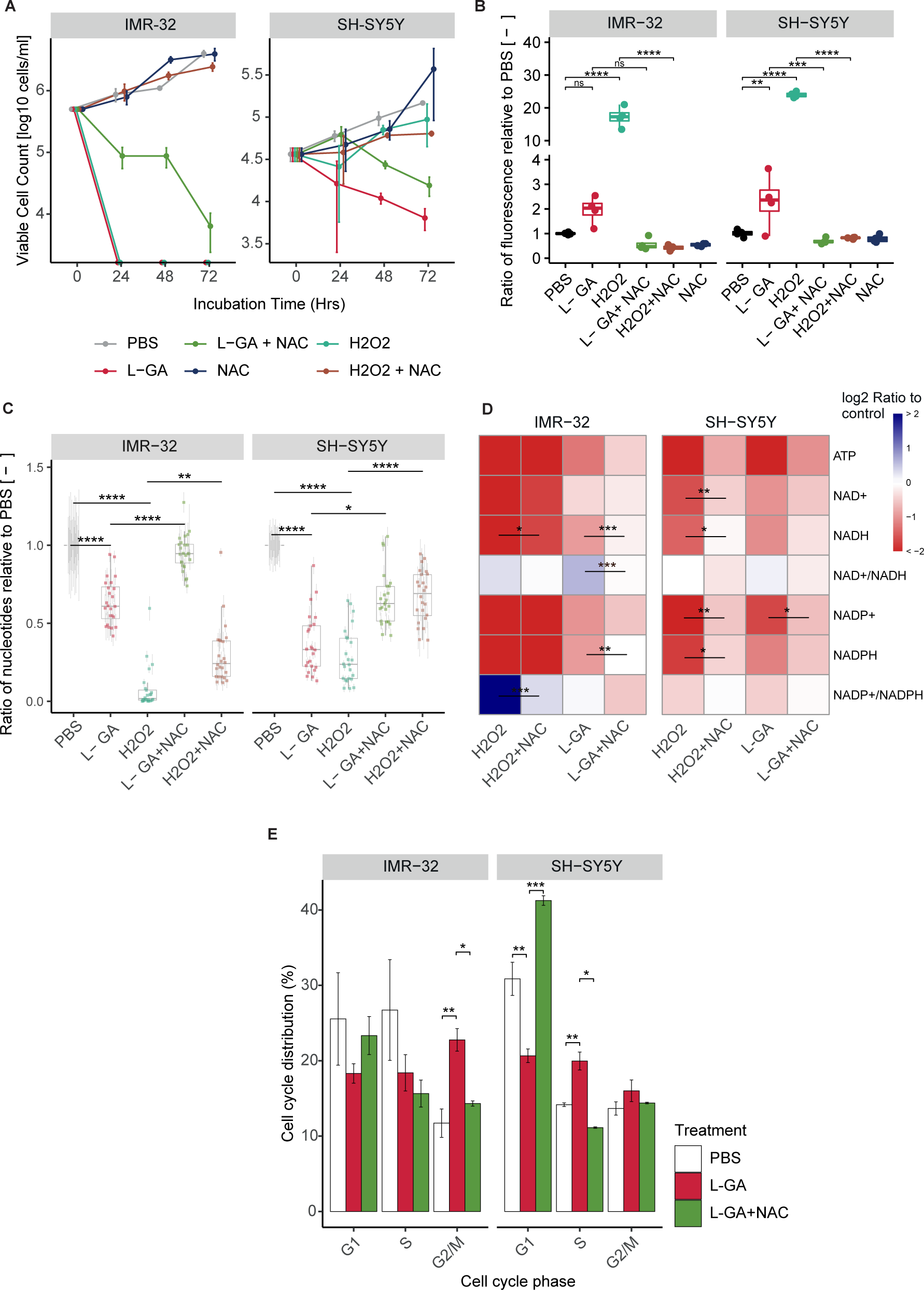
L-GA induces oxidative stress which is linked to cell growth, nucleotide synthesis and the cell cycle. **A** IMR-32 and SH-SY5Y were treated for 72 hrs with 1 mM L-GA, 100 µM H_2_O_2_ and 5 mM NAC, in the combinations shown. Viable cell counts were taken every 24 hrs. Error bars represent the *±* SEM (n=3). **B** IMR-32 and SH-SY5Y were treated for 4 hrs in combinations as in A. ROS species were quantified by measuring the fluorescence from DCFDA. Box plots show the ratio of fluorescence relative to the PBS control (n=4). P-values were calculated using Anova and Tukey post-hoc analysis. **C** IMR-32 and SH-SY5Y were treated with drug combinations as in A. Nucleotides were extracted and measured by DI-MS/MS. Boxplots present the mean ratio of nucleotide intensity normalised to PBS, error bars represent *±* SEM (n=4). P-values were calculated using an Anova and Tukey post-hoc analysis. **D** Cells were treated as in C; intensities of ATP, NAD+, NADH, NADP+ and NADH with associated ratios were calculated and normalised to PBS (n=4). P-values were calculated using Anova and Tukey post-hoc analysis. **E** Cell cycle analysis was performed in IMR-32 and SH-SY5Y cells following treatment for 24 hrs with PBS, 1 mM L-GA or 1 mM L-GA + 5 mM NAC. Cells were gated into phases for Anova and Tukey post-hoc analysis on the cell cycle distributions (n=3).

ROS generation can be monitored by the fluorometric analysis of the oxidation of 2’,7’ –dichloroflu-orescin diacetate (DCFDA) by ROS into 2’, 7’–dichlorofluorescein (DCF). We assessed whether L-GA induces ROS generation and if this can be rescued by NAC. Cell lines IMR-32 and SH-SY5Y were treated with: 1X PBS, 1 mM L-GA, 100 µM H_2_O_2_, 1 mM L-GA + 5 mM NAC, 100 µM H_2_O_2_+ 5 mM NAC and 1X PBS + 5 mM NAC. Cells were incubated for 4 hrs with each treatment condition before DCFDA staining and fluorescence reading, Figure 5B shows fluorescence intensity after treatment normalised to PBS treated cells. In both cell lines 100 µM H_2_O_2_, induced a significant increase in fluorescence in IMR-32 and SH-SY5Y (*p ≤* 0.001). L-GA induces an increase in fluorescence in both cell lines. Upon treatment with 5 mM NAC, the fluorescence of 100 µM H_2_O_2_treated cells decreased significantly (*p ≤* 0.01). Similarly, there is a reduction in fluorescence of L-GA treated cells in the presence of NAC. From this data, we concluded that 1 mM L-GA induces an increase in ROS. As we had previously found L-GA to deplete nucleotides, we questioned as to whether nucleotide pools are sensitive to oxidative stress and if NAC could recover depletion.

Nucleotides were measured after 24 hrs treatment with 1 mM L-GA, 100 µM H_2_O_2_ and 5 mM NAC and in the combinations shown in Figure 5C. As found previously, there was a significant reduction in nucleotide pools following L-GA treatment (*p ≤* 0.0001). Upon 100 µM H_2_O_2_ treatment, nucleotides were significantly reduced in both cell lines (*p ≤* 0.0001). The co-application of L-GA and NAC results in a significant increase in nucleotide pools relative to L-GA alone (*p ≤* 0.05). The application of NAC to 100 µM H_2_O_2_ treated cells also significantly increased the nucleotide pools, relative to 100 µM H_2_O_2_alone (*p ≤* 0.01).

We examined ATP, NAD+, NADH, NADP+ and NAPDH with associated ratios, following ROS treatment (Figure 5D). The ATP concentration was reduced significantly upon L-GA and H_2_O_2_ treatment (*p ≤* 0.01) in both cell lines. The addition of NAC partially remedied L-GA and H_2_O_2_ dependent depletion of ATP. L-GA caused a decrease in NAD+ and NADH with an increase in the NAD+/NADH ratio in both cell lines. We found that, in IMR-32 cells, 100 µM H_2_O_2_ caused a slight increase in the NAD+/NADH ratio. The addition of NAC to L-GA and H_2_O_2_ treated cells reduces the NAD+/NADH ratio, with the strongest increase in NADH levels. Ultimately availability of NAD(P)+ and NAD(P)H is compromised in both cell lines. Although nucleotide pools- and ATP in particular-were depleted in both cell lines by H_2_O_2_, growth of SH-SY5Y cells persisted. This suggests that induction of ROS production and nucleotide depletion, are not fully responsible for cell growth inhibition by L-GA. Nucleotide depletion is likely to be a concerted phenotype in the multi-modal action of L-GA.

Following the findings of partial recovery of nucleotide pools via NAC, we hypothesised that the cell cycle block induced by L-GA could be recovered by NAC. IMR-32 cells and SH-SY5Y cells were challenged with 1 mM L-GA or 1 mM L-GA + 5 mM NAC for 24 hrs (Figure 5E). Cells were harvested and stained with propidium iodide and analysed by flow cytometry. As found previously, L-GA caused a significant increase in the G2/M phase (*p ≤* 0.01) in IMR-32 cells. The co-application of NAC significantly relieved the G2/M block relative to L-GA alone (*p ≤* 0.05). In SY-SY5Y cells L-GA caused a significant decrease in the G1 phase and a significant increase in the S phase (*p ≤* 0.01). Upon co-application with NAC we found a significant increase in the G1 phase (*p ≤* 0.001) and a significant decrease in the S phase (*p ≤* 0.05) relative to L-GA alone.

In summary, L-GA induces ROS generation, dysregulates NAD(P)+ and NAD(P)H availability, depletes nucleotides and causes cell cycle arrest. The co-application of NAC partially relieves these cell states induced by L-GA. NAD+ and NADH generation is tightly linked to central carbon metabolic processes. We aimed to assess whether NAC mitigates L-GA activity as a glycolytic inhibitor.

### N-acetyl cysteine partially restores glycolytic function of L-GA treated cells

Cell lines IMR-32 and SH-SY5Y were treated for 1hr or 16 hrs with 1X PBS, 1 mM L-GA or 1 mM L-GA + 5 mM NAC. Within the treatment time cell culture media was exchanged supplemented with U-^13^C-glucose to initiate metabolite labelling. Labelling was quenched at 10 mins, metabolites were extracted and subjected to GC-MS (Figure 6).

**Figure 6.**
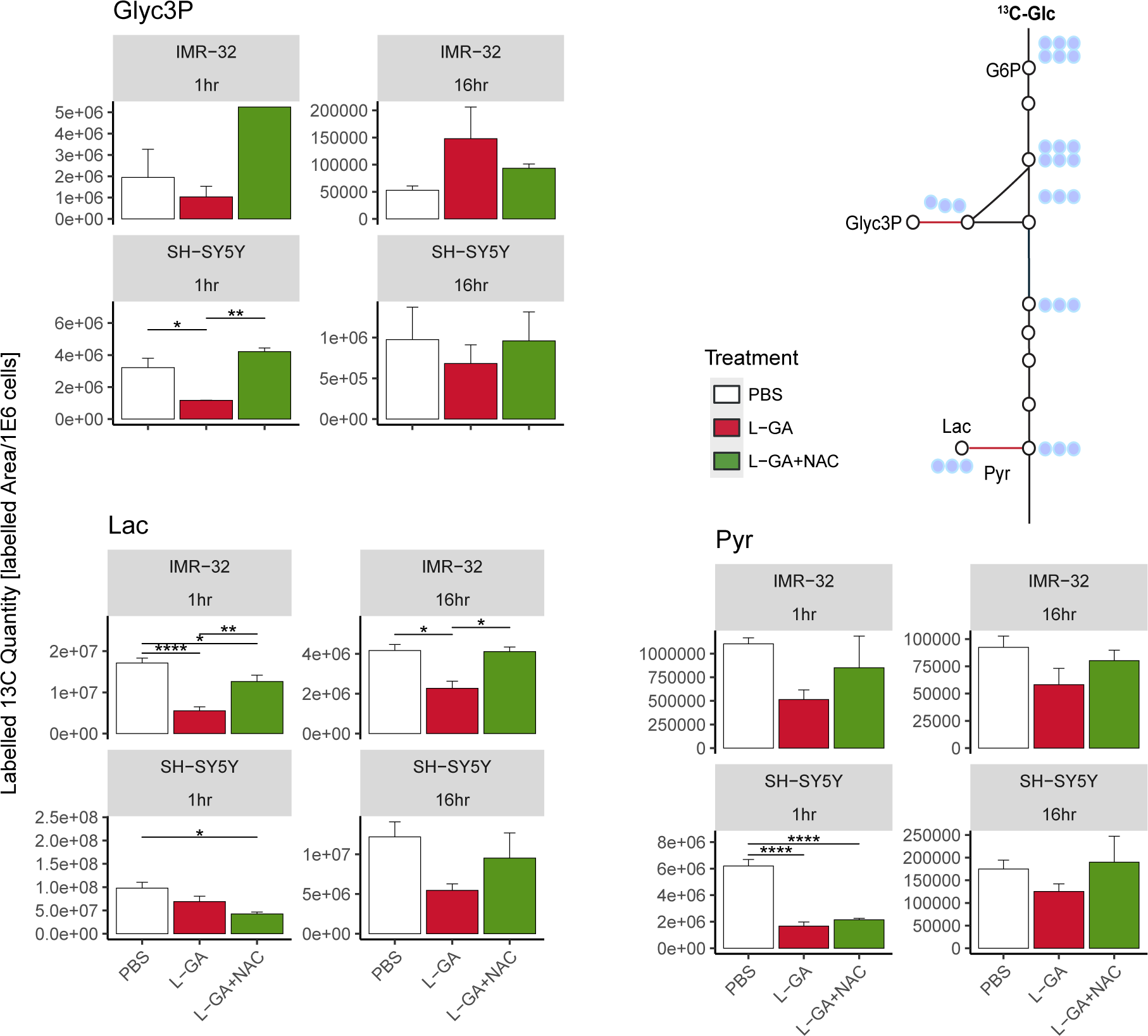
The antioxidant N-acetyl-cysteine partially restores metabolic function of L-GA treated neuroblastoma cells. Polar metabolites were extracted from IMR-32 and SH-SY5Y cell lines following treatment of 1 mM L-GA or 1 mM L-GA + 5 mM NAC for 1 and 16 hrs and 10 mins labelling with U-^13^C-glucose. Samples were processed via GC-MS and analysed to attain the peak area and label incorporation to calculate labelled area for each metabolite. Bar charts depict the mean labelled ^13^C quantity (Labelled area/1E6 cells) (m/z +3). Error bars represent ± SEM, (n=3-4, except IMR-32: Glyc3P, 1hr L-GA+NAC (n=1)). P-values were calculated using Anova and Tukey post-hoc test.

For SH-SY5Y cells, we found a decrease in ^13^C-labelled quantities of Gly3P, Lac and Pyr following 1 hr treatment with L-GA. Co-application of L-GA+NAC significantly increased ^13^C-labelled quantities in Glyc3P, however no effect was observed with Lac or Pyr. After 16 hrs treatment, Lac showed a stronger response to L-GA treatment which was mitigated with NAC co-application. Labelled quantities of Glyc3P and Pyr were slightly reduced upon L-GA application relative to PBS, with L-GA+NAC showing marginal increased labelled quantity relative to L-GA alone.

For L-GA treated IMR-32 ^13^C-labelled quantities for Glyc3P, Pyr and Lac were decreased after 1 hr treatment. The addition of NAC at this time-point partially mitigated L-GA dependent inhibition at Pyr and Lac. After 16 hrs treatment, Pyr and Lac labelled quantities remained lower than the control for L-GA cells. The addition of NAC partially restored labelled quantities in a similar fashion to 1 hr with Pyr and Lac.

In summary, we find that NAC partially recovers L-GA mediated inhibition of glycolysis, particu-larly at Lactate. We suspect this to be a result of the NAD+/NADH cycling governed by glyceraldehyde-3-phosphate dehydrogenase (GAPDH), lactate dehydrogenase (LDH) and glycerol-3-phosphate dehy-drogenase (GPD). The sensitivity of GAPDH to oxidative stress is well documented, although LDH and GPDH are not known to be ROS sensitive, however both enzymes require NADH, so with limited NADH availability their function decreases. We observed a heterogeneous response to L-GA between cell lines, particularly at Glyc3P. IMR-32 appeared to respond to NAC mediated recovery of glycolytic much earlier than SH-SY5Y, this may be due to the difference in rate of L-GA metabolism in each cell line. It is likely that L-GA acts as a direct inhibitor of glycolysis and secondly as a ROS-generator with latent indirect effects which are partially recovered by NAC. To further address this bi-modal effect we examined post-translational modification to the proteome.

### Phosphoproteomic analysis reveals cell cycle dependent kinases are perturbed by L-GA

A phosphoproteomic approach was performed in order to derive information about which signalling pathways are modulated by L-GA. In addition, we aimed to show that affected pathways are relieved via NAC application. IMR-32 cells were treated with 1X PBS, 1 mM L-GA or 1 mM L-GA + 5 mM NAC for 4 hours before harvesting. Enriched phosphoproteomic samples were measured via LC-MS. Intensities of peptide sequences with phosphorylation sites were annotated with associated gene names and kinases using the PhosphoSitePlus database (32). The fold change in phosphopeptide intensities (L-GA/PBS) were calculated, mapped to the parent protein and plotted against the -log10(p-value) (Figure 7A). Phosphopetides were also annotated with predicted kinases that phosphorylate the peptide. Phosphopetides phosphorylated by kinases: ERK2, AuraA;CDK1, CK2A1 GSK3A, MAPKAPK2 were found to be the top 5 regulated following L-GA treatment. When assessing the proteins mapped to the phosphopetide, we found the strongest response from PARVA, TP53, DTD1, MAPT and CDC25B. Although not annotated with an associated kinase we found strong up-regulation of ATN1 and down-regulation of CRAMP1L.

**Figure 7.**
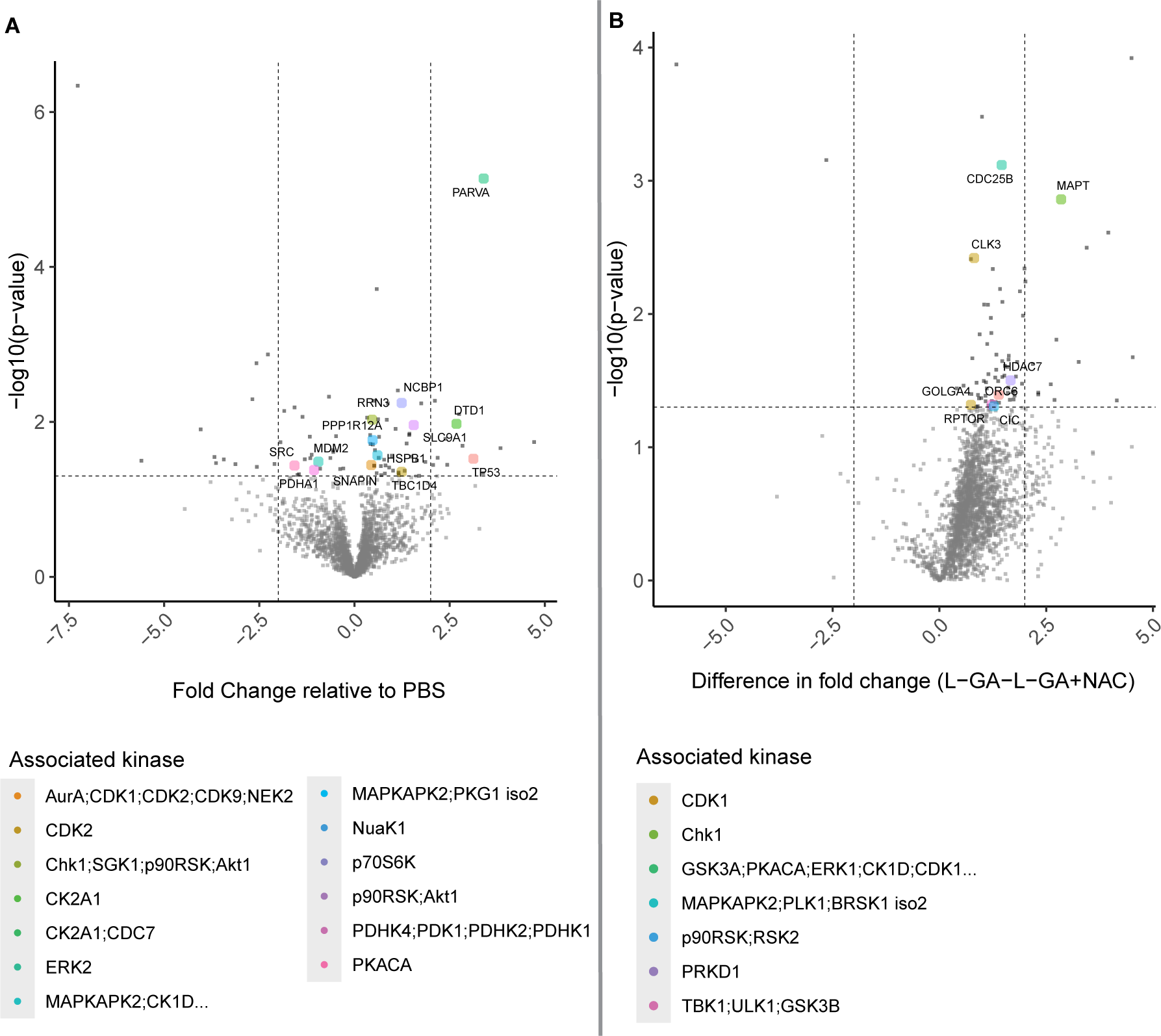
Phosphoproteomic analysis of L-GA treated cells. **A** IMR-32 cells were treated for 4 hrs with 1 mM L-GA, 1 mM L-GA+NAC and PBS and harvested for phosphoproteomic analysis (n=4). Phosphopeptides were measured by LC-MS and fold change changes were calculated for L-GA/PBS and L-GA+NAC/PBS. Significant phosphopeptides were found by Anova followed by Tukey post-hoc tests, coloured by the associated kinase. Phosphopeptide sequences were annotated with a protein name and associated kinase. **B** Fold change data between L-GA and L-GA+NAC treatment were passed to a two-tailed T-test. Significant peptides were annotated with protein names and associated kinases and highlighted in red (n=4).

**Figure 8.**
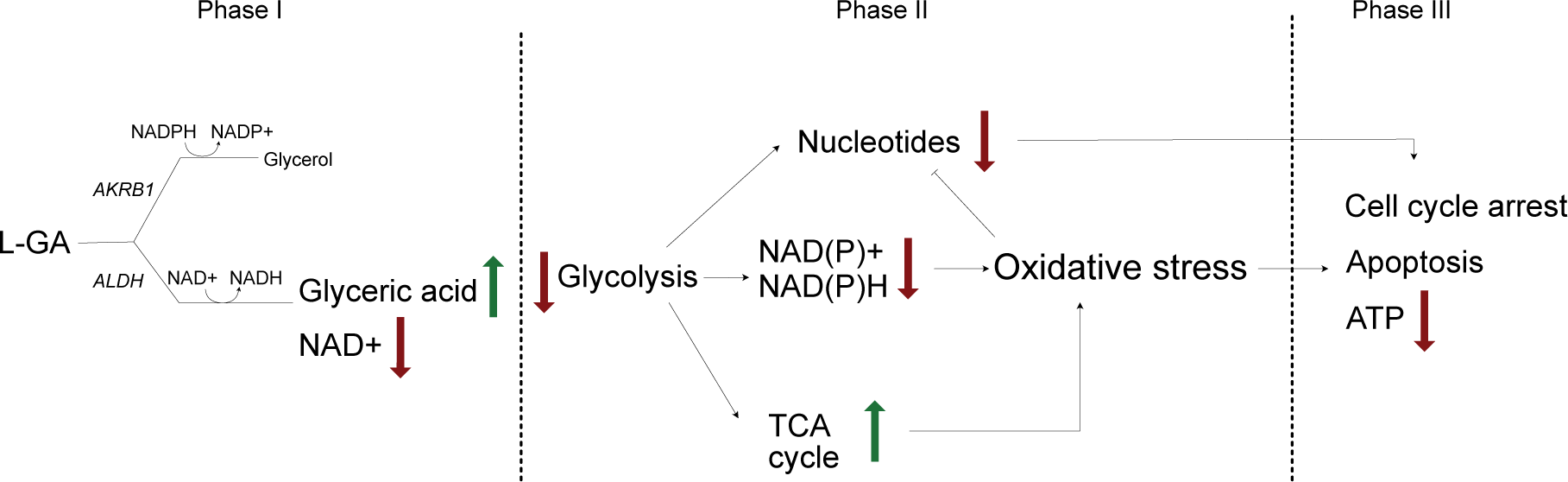
Proposed mechanism of L-GA on cellular metabolism. L-GA is rapidly converted to glyceric acid, favoured over glycerol metabolism, consuming NAD+. At adequate concentrations of L-GA, glycolysis is inhibited resulting in further decrease of ATP and NAD derived co-factors. This results in an increase of oxidative stress which co-currently up-regulates the TCA cycle, leading to ROS production and further oxidative stress. Nucleotides are depleted as a facet of cell stress and glycolysis is further inhibited. As a sum of parts, the cell enters cell cycle arrest and initiates apoptosis.

In order to discern which phoshopeptides were most significantly affected by the addition of NAC, we examined the relative difference on phoshopeptide intensities between L-GA and L-GA+NAC treatment (Figure 7B). We found the largest difference in kinases: GSK3A;PKACA;ERK1;CK1D;CDK2. Therefore, phosphopeptides regulated by these kinases, modulated by L-GA, were relieved by NAC. Although not annotated with a kinase we found NCOA3, RANGAP1 and GTPBP1 to be the strongest responders to NAC following L-GA treatment.

We find that L-GA exhibits a post-translational effect on cells at the phosphoproteomic level. The addition of NAC appears to relieve pathways modulated by L-GA. Interestingly, the top modulated pathway, ERK2, has a broad effect on cellular metabolism, cell cycle regulation and cell adhesion. This modulation correlates with the the cell cycle analysis, actin staining and metabolomics experiments performed previously.

## Discussion

Homeostasis is essential for cancer cell survival. We have shown that by challenging the cell with - a long-time presumed - glycolytic inhibitor, homeostasis is severely affected. Subsequently, a novel mechanism is presented for L-GA action, in which multiple facets of cancer cell metabolism are dysregulated and not limited to glycolytic inhibition. Analysis of nucleotide metabolism, cell cycle arrest and (phospho)proteomics presented L-GA induced oxidative stress. This mode of action of L-GA is previously unreported in neuroblastoma cells, and is of particular significance in understanding its therapeutic potential.

The function of L-GA as a glycolytic inhibitor has dominated the focus of its previous research, however the mechanism was not characterised. Employing the pSIRM strategy showed that L-GA decreases the carbon flow into metabolites that require NAD(H) as a co-factor. Through the integration of GC- and DI-mass spectrometry data sets, it was indicated that L-GA breaks the cycling of cytosolic NAD+/NADH. This breakage may impact the redox state of the cell thereby inhibiting metabolism. Glyceraldehyde dehydrogenase (GAPDH) produces cytosolic NADH via the metabolism of glyceraldehyde-3-phosphate to 1,3-glyceric acid biphosphate. GAPDH possess a cysteine rich active site, thereby rendering it sensitive to ROS (33). We suspect that GAPDH activity may be impacted by L-GA directly by delivering smaller amount of the substrate glyceraldehyde-3-phosphate, and indirectly as elevated ROS levels via L-GA inhibit GAPDH. D,L-glyceraldehyde has been shown to: raise intracellular peroxide levels, dysregulate NAD(P)/NAD(P)H ratios, acidify the cell and inhibit glycolysis - at more than double the concentration used in this paper - in islet cells via non-mitochondrial pathways. This phenomenon causes oxidative stress which is remedied with NAC application (20, 34). Hydrogen peroxide has been shown to inhibit GAPDH activity. This results in reducing carbon flow into lower glycolysis intermediates and upper glycolysis intermediates accumulate (35). The reduction of carbon flow complements our data with respect to L-GA, although Van Der Reest et al (2018), reported depletion of pyruvate levels where we observed pyruvate accumulation in the time frames employed. Furthermore, we found that D-GA has a lesser effect on label incorporation into Lactate and Glycerol-3-phosphate than L-GA, nor does it reduce label incorporation into glyceric acid-3-phosphate. D-GA may be metabolised by ALDO(A/C) to produce fructose-1-phosphate which can be used to re-fuel glycolysis, L-GA may produce sorbose-1-phosphate, which inhibits hexokinase (36). L-GA did not cause a notable accumulation of upper glycolysis intermediates, as would be expected with GAPDH inhibition, which may be a result of hexokinase inhibition. We found that ALDOC protein levels in SH-SY5Y to be higher than IMR-32, which may give reasoning to the lower response to D-GA (supplementary Figures 9I). This was complemented by nucleotide analysis in which, D-GA did not show the same level of nucleotide depletion as caused by L-GA in IMR-32.

We found that IMR-32 and SH-SY5Y respond to L-GA differently, particularly in the early stages of L-GA treatment, it appears that SH-SY5Y is more sensitive to NADPH depletion, as well as nucleotide deletion. This may be due to higher levels of ARKB1 which consumes NAPDH in the production of glycerol from L-GA (supplementary Figures 9I). IMR-32 tended to be more sensitive to glycolytic inhibition and NAD(H) depletion. This may be due to the metabolic rates of the cells and their relative dependency on subsequent metabolic pathways. Specifically, we found varying levels of aldehyde dehydrogenase and aldo-keto reductase between cell lines (supplementary Figures 9I). This is certainly an issue of consideration when discussing L-GA efficacy and mechanism between cell types.

The application of L-GA and H_2_O_2_ to neuroblastoma cells induced a depletion of nucleotide pools. Literature on the effect of ROS on the nucleotide intermediate metabolism is sparse. Although much is known about the effect of ROS on depletion of ATP production and DNA damage, the work here provides novel insights to how ROS affects nucleotide intermediates. The application of NAC in the presence of H_2_O_2_ and L-GA indicates that nucleotide synthesis is indeed ROS sensitive, although may not be necessary for cell survival, as in the case of sustained SH-SY5Y cell growth following H_2_O_2_ application. We suspect that L-GA-derived ROS and nucleotide depletion alone does not induce apoptosis but rather the sum of all L-GA modes of action result in cell death. Despite this, further work is required to identify the specific enzymes which are most susceptible to ROS. It is unclear whether it is a direct action of ROS on nucleotide metabolising enzymes or whether it is a result of signalling pathway perturbation. Enzymes associated with nucleotide biosynthesis: RNR and TYMS, are known to be ROS sensitive (37, 38). However, our results suggest that additional nucleotide biosynthesis enzymes are subject to modulation via ROS, directly or indirectly. Through the measurement of the product/substrate ratios of nucleotide intermediates it was evident that SH-SY5Y and IMR-32 respond differently to oxidative stress. Curiously, with early intermediates such as SAICAR displayed accumulation upon ROS exposure (supplementary Figures 9F). Characterisation of all nucleotide intermediates via DI-MS/MS is expected to precisely reveal additional L-GA targets within the nucleotide biosynthesis pathway.

Albonzinia et el. (2022) showed that MYCN amplified neuroblastoma cells exhibit a high demand for cysteine (39). MYCN amplification results in the over expression of anti-oxidant genes to maintain the redox balance, which is stressed due to depleted cysteine from high MYCN levels. MYCN levels in IMR-32 are amplified (100 MYCN copies) compared to a gain (3 MYCN copies) in SH-SY5Y. Given that our observations showed IMR-32 to be more sensitive than SH-SY5Y to H_2_O_2_, it is possible that the basal state of IMR-32 is under more oxidative stress due to the amplified MYCN.

Phosophoproteomic data highlighted kinase pathways which were modulated via L-GA treatment. Specifically, GSK3*β* and MAPKAPK2 axes identified by MAPT and PARVA. MAPKAP2 is known to regulate the G2/M transition in response to DNA damage via Cdc25B/C phosphorylation (40). Given that L-GA depletes nucleotides, it is likely that DNA damage ensues and appropriate responses are initiated. Indeed, we found Cdc25B/C phosphorylation to be up-regulated upon L-GA treatment, giving credence to MAPKAPK2-14-3-3 mediated cell cycle arrest and apoptosis (supplementary Figures 9G). Furthermore, we found p53 associated with AurA and CDK1 kinases to be up-regulated in response to L-GA, which is indicative of DNA damage response and G2/M arrest (41, 42). AurA has been implicated in high-risk neuroblastoma and there are currents efforts to validate AurA inhibitors (43). It would be of clinical interest to examine the efficacy of AurA inhibitors and L-GA co-application in neuroblastoma models.

Many chemotherapeutics induce ROS, pushing the cancer cell to lethal limits (44). Doxorubicin, Daunorubicin and cisplatin induce high levels of ROS (45, 46). Nucleotide analogs such as 5-fluoropyrimidine produce lower levels of ROS (47). L-GA acts in a similar manner however possesses significantly lower toxicity. In a complementary context, our data are supported by the findings of Uetaki et.al. The authors showed that high levels of ascorbic acid induces strikingly similar effects in MCF7 human breast adenocarcinoma and HT29 human colon cancer cells as L-GA does in neuroblastoma cells. Ascorbic acid induces HMOX1 over expression, depletes dNTPs and inhibits glycolysis by NAD+/NADH dysregulation. However, global depletion of nucleotides and their intermediates via ascorbic acid was not shown (48).

The targeting of cancer cell metabolism by the glycolytic inhibitors 2-DG and BrPyr have shown to be ineffective or toxic due to narrow therapeutic windows (49, 50, 51, 52). Yet in light of their molecular mechanism, BrPyr has been shown to induce ROS increase and activation of GSK3*β*, as found with L-GA treatment (53, 54, 55). In combination with MAPKAPK2 and TP53 up-regulation, glycolytic inhibitors are shown to induce DNA damage responses and cell cycle arrest.

Despite 80 years of research L-GA has yet to take hold as a viable therapeutic. However, L-GA is of particular clinical interest due to its low toxicity, being relatively inexpensive and easy to synthesise. When bracketed in a manner similar to chemotherapeutics, the therapeutic concentration of L-GA required is relatively high (56, 49, 57). We found that the IC50 of 5 neuroblastoma cell lines to be in the range of 262-1005 µM after 24 hrs treatment, with nucleotide depletion occurring above 0.5mM L-GA (supplementary Figures 9B). However, this is only when considering cell culture models and has not yet been shown to translate to *in vivo* trials. The clearance of GA from the body is rapid. Brock and Niekamp found that a working concentration of 2 mM GA could not be maintained in rabbits (58). The application of the compound closer to the therapeutic target site via peritoneal injection presented promising results (52). In addition, our data suggests that maintenance of a working concentration may not be necessary as we found that L-GA treated cells could not recover after L-GA was removed.

In the context of neuroblastoma, the conventional regimen of vincristine [O], cisplatin [P], etoposide [E], and cyclophosphamide [C], OPEC, dosages are in the micromolar range. Yet, patients experience severe side effects ranging from hair loss, appetite suppression, nausea and increased chance of resistance (59). Due to the high molarity of L-GA required for potential therapy, it is unlikely to translate into a mono-therapy. However, L-GA has scope to be used as an adjunct to provide more efficacy to OPEC regimens, reducing the concentrations and time for chemotherapy treatment. Indeed, 2-DG and DL-GA act in synergy to reduce tumour growth of ehrlich ascites carcinoma (52), however translation to neuroblastoma treatment regimens is to be explored.

---

**Figure 9.**
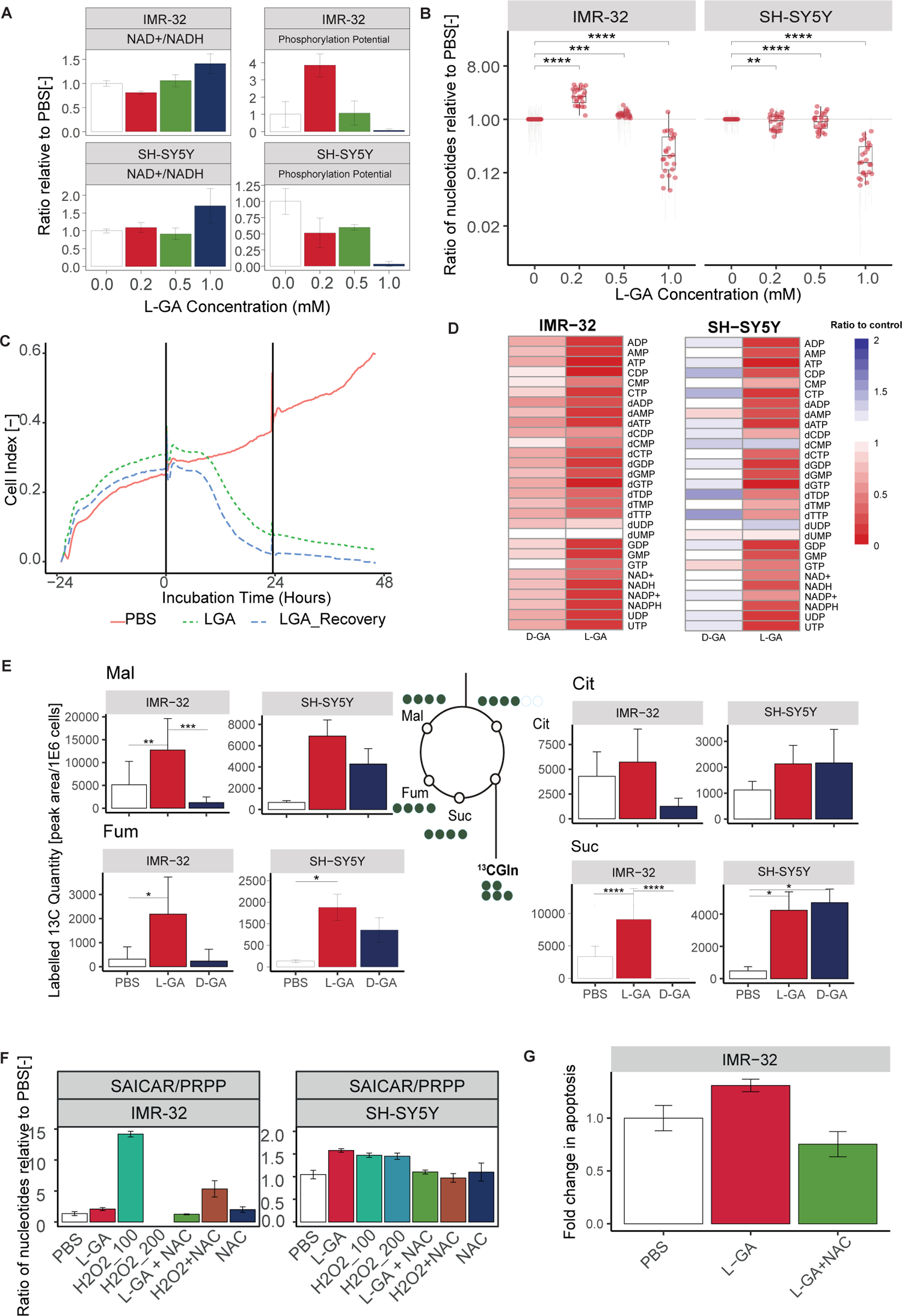
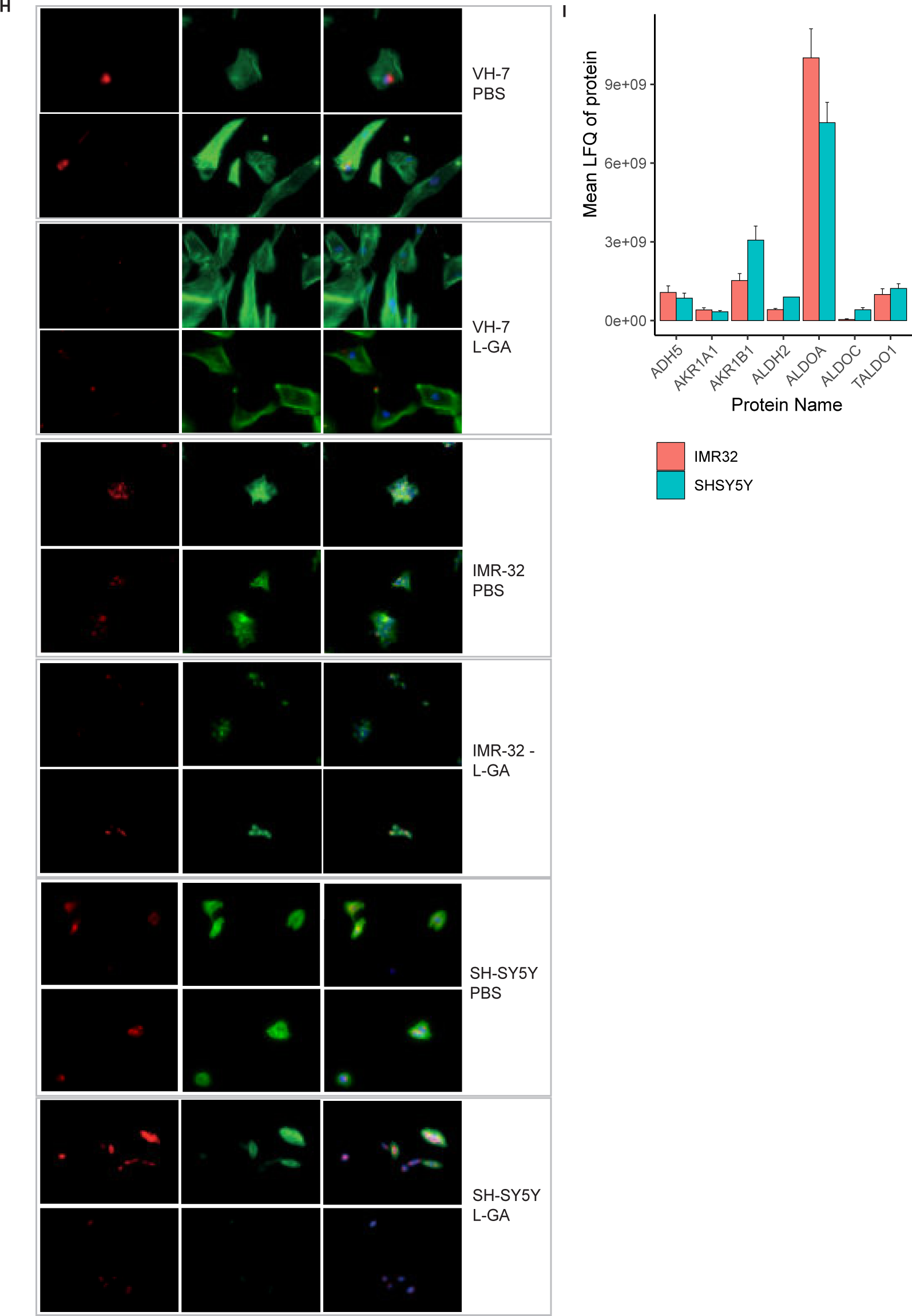
Supplementary figures. **A** IMR-32 and SH-SY5Y cells were treated with a range of L-GA concentrations for 24 hrs, nucleotide intensities were measured by DI-MS and used to calculate the mean NAD/NADH and phosphorylation potential ratio and normalised to PBS, errors bars show *±* SEM (n=3). **B** IMR-32 and SH-SY5Y cells were treated with a range of L-GA concentrations for 24 hrs, DNA and RNA nucleotide intensities were measured by DI-MS and normalised to PBS (n=3). **C** IMR-32 cells were seeded in an iCelligence device and allowed to settle before exchanging of media for 1 mM L-GA or PBS. Following 24 hrs incubation media was exchanged with fresh media or media containing 1 mM PBS to simulate recovery. **D** IMR-32 cells were treated with 1 mM L-GA or 1 mM D-GA for 24 hrs, nucleotide intensities were measured by DI-MS and normalised to PBS for heatmap representation (n 3). **E** Cell lines IMR-32 and SH-SY5Y were treated with 1 mM L-GA or 1 mM D-GA for 1 hr and labelled for 30 mins with U-^13^C-glutamine within the treatment time. Bar charts depict the mean labelled ^13^C-quantity (Labelled peak area/1E6 cells) of depicted metabolites (m+4). Error bars represent *±* SEM (n=3) P-values were calculated using Anova and Tukey post-hoc test. **F** IMR-32 and SH-SY5Y were treated for 24 hrs with 1 mM L-GA, 100 µM H_2_O_2_ and 5 mM NAC and in the combinations shown. Nucleotides were extracted from cell lysates and measured by DI-MS/MS. Bar chart present the mean intensity of product substrate ratios normalised to PBS, error bars *±* SE (n=4). **G** IMR-32 cells were incubated for 5 hours with 1 mM L-GA, 1 mM L-GA + 5mM NAC. For each replicate 10.000 cells were measured by flow cytometry. Bar charts represent the mean *±* SEM (n=3). **H** Cell lines VH-7, IMR-32 and SH-SY5Y were fixed to glass slides and stained with Phallodin-iFluor(F-) or Deoxyribonuclease I (G-) actin probes then imaged at 40X magnification using Keyence Bz-x700 microscope. **I** Proteomic analysis of cell lines: SH-SY5Y and IMR-32. Data represents the mean label free quantity of untreated cells, error bars represent *±* SEM (n=3).

**Table 1.** IC50 of a panel of neuroblastoma cell lines treated with L-GA via WST assay over 96 hours treatment time.

| Cell line | MYCN-Status | IC50 24 | IC50 48 | IC50 72 | IC50 96 | Delta IC50 |
| --- | --- | --- | --- | --- | --- | --- |
| BE(2)-C | amplified | 1005 | 285 | 137 | 447 | 469 |
| SH-SY5Y | gain | 660 | 783 | 94 | 300 | 460 |
| SK-N-AS | single copy | 1001 | 622 | 228 | 1069 | 730 |
| CLB-GA | single copy | 995 | 98 | 110 | 103 | 104 |
| GI-M-EN | single copy | 262 | 196 | 110 | 214 | 196 |
| VH-7 | - | 1042 | 963 | 301 | 244 | 638 |

## Methods

### Cell Culture

Cell lines were maintained in DMEM (Gibco) medium, without glucose, glutamine, phenol red and sodium pyruvate. Media was supplemented with 10% fetal bovine serum (Gibco), 2 mM glutamine (Thermo Fisher), and 2.5 g/L glucose (Merck) and cultivated at 37 *^◦^*C, 5% CO_2_, 21% O_2_ and 85% relative humidity. To avoid contact inhibition, cells were passaged every 3-4 days. The neuroblastoma cell lines: BE(2)-C (RRID: CVCL_0529) was obtained from ECACC (Salisbury, UK). IMR-32 (RRID: CV CL_0346), GI-ME-N (RRID: CVCL_1232) and SH-SY5Y (RRID: CVCL_0019) cell lines from the DSMZ (Braunschweig, Germany). Cell lines were authenticated via the Multiplex human Cell line Authentication Test (Multiplexion, Immenstaad, Germany). The active primary human foreskin fibroblasts from an infant donor (VH7) were a gift from Petra Boukamp (German Cancer Research Center (DKFZ), Heidelberg, Germany)

### Proliferation assay

Numbers of viable cells were determined using trypan blue exclusion. A cell suspension was prepared with 50 µL 0.4% trypan blue solution (w/v) and 50 µL trypsinised cells. Following mixing 10 µL of the cell suspension was loaded into a cell counting slide. Cell numbers were measured using the TC10 automated cell counter (Biorad).

### Proteomic Analysis

Cells were harvested by scratching cells in 2 ml ice-cold 1X PBS on ice and transferred to a 2 ml tube, cells were centrifuged at 14,000 x *g* for 5 mins, 4 *^◦^*C. Cell pellets were lysed in 300 µL urea buffer (8 M Urea, 100 mM TrisHCl (pH 8.5). Lysates were sonicated for 30s and centrifuged at 14,000 x *g* for 5 mins, 4 *^◦^*C. Lysates were then incubated for 10 mins at 4 *^◦^*C under constant agitation. Lysates were centrifuged at 14,000 x *g* for 5 mins, 4 *^◦^*C, supernatants removed and protein concentration determined via the Pierce BCA Protein Assay Kit (Thermo Fisher). The absorbance of each sample was read at a wavelength of 562 nm (Infinite 2000, Tecan). Each sample was measured in technical duplicates.

Proteins were alkylated and denatured by addition of 2 mM DTT for 30 min at 25 *^◦^*C, followed by addition of 11 mM iodoacetamide for 15 min at room temperature in the dark. One hundred µg of protein was aliquoted and digested using Lys-C (Wako, 1:40 (w/w) and immobilised trypsin beads (Applied Biosystem, 5-10 µL, 4 hrs under rotation, 30 *^◦^*C). Digested proteins were diluted with 50 mM ammonium bicarbonate before tryptic digestion. Digestion was halted with 5 µL of triflouracetic acid (TFA). Twenty µg of digested sample was then desalted and purified on in-house prepared stage tips. Stages tips were primed with 50 µL 100% methanol and 50 µL 0.5% acetic acid (v/v). Stage tips were then centrifuged at 300 x *g* for 7 mins and digests were loaded into the tips and washed once with 50 µL 0.5% acetic acid (v/v). Peptides were eluted by addition of 10µL 0.5% acetic acid (v/v) in 80% acetonitrile. Eluates were dried by centrifugation under vacuum then resuspended in 10 µL 0.5% acetic acid (v/v) and sonicated at room temperature for 5 mins.

For phosphoproteomic analysis, digested samples were desalted using SepPak C18 columns prior to phosphopeptide enrichment using the High-Select TiO2 Phosphopeptide Enrichment Kit (Thermo Scientific). Samples were resuspended in 150µL of Binding/Equilibration Buffer and vortexed. Samples were then enriched for phosphopeptides using a TiO2 Spin Tip, washed and eluted in preparation for LC-MS.

### Analysis of peptides via liquid-chromatography mass spectrometry (LC-MS)

Peptide mixtures were analysed by LC-MS following a shotgun proteomics method (60). LC-MS was performed on an automated high performance liquid-chromatograph (NanoLC 415, Eksigent) coupled to tandem mass spectrometry (Q Exactive HF, Thermo Fisher). Samples were pipetted into 5 µL aliquots into a 96-well plate to be acquired in two technical replicates. Samples were loaded on the column with a flow rate of 450 nL/min. Elution was performed with a flow rate of 400 nl/min using a 240 mins gradient ranging from 5% to 40% of buffer B (80% acetonitrile and 0.1% formic acid) in buffer A (5% acetonitrile in 0.1% formic acid). Chromatographic separation was performed on a 200 cm long MonoCap C18 High Resolution 2000 column (GL Scientific). The nanospray source of the Q Exactive HF was maintained at 2.4 kV and the downstream ion transfer tube at 260 *^◦^*C. Data were acquired in data-dependent mode with one survey MS scan (resolution: R=120,000 at m/z 200) followed by a maximum of ten MS/MS scans (resolution: R=30,000 at m/z 200, intensity threshold: 5,000) of the most intense ions.

Raw data were analyzed using the MaxQuant proteomics pipeline (version 1.5.3.30) and the built in Andromeda search engine with the human Uniprot database (61). Carbamidomethylation was set as a fixed modification, oxidation of methionine as well as acetylation of N-terminus as variable modifications. The search engine peptide assignments were filtered at 1% FDR. Peptides with a minimum length of seven amino acids and a maximum of two miscleavages were further processed. Peptides were matched between runs. Label-free quantification (LFQ) was performed for all samples, using razor peptides to define groups of peptides and unique proteins belonging to the razor peptides groups with the largest matches. The software Perseus (version 1.5.6.0) was used to collate LFQ data, perform imputation and log_2_ transformation of data prior to input of data into custom R script for further analysis using version 4.0.4 (62).

The mass spectrometry proteomics data have been deposited to the ProteomeXchange Consortium via the PRIDE (63) partner repository with the dataset identifier PXD048695.

### Metabolomics

Medium was changed 24 and 4 hours before harvesting cells for metabolome analysis using pulsed stable isotope-resolved metabolomics (64) in tandem with absolute quantitative gas chromatography (GC-MS). Cells were labelled prior to harvest with medium containing either 14 mM U-^13^C-glucose or 14 mM ^12^C-glucose for glucose tracing and 2 mM U-^13^C-glutamine or 2 mM ^12^C-glutamine for glutamine tracing. Harvested cells were washed with HEPES buffer (140 mM NaCl, 5 mM HEPES, pH 7.4) containing labeled or unlabeled glucose/glutamine and quenched by adding 50% ice-cold methanol containing 2 *µ*g/mL cinnamic acid (Merck) as an internal control. Polar metabolites were extracted as previously described (65). Metabolite analysis was performed on a 1D gas chromatograph coupled to time of flight mass spectrometer (GC-ToF-MS, Pegasus IV-ToF-MS-System, LECO), samples were handled with an auto-sampler (MultiPurpose Sampler 2 XL, Gerstel). The samples were injected in split mode (split 1:5, injection volume 1 µL) in a temperature-controlled injector (CAS4, Gerstel) with a baffled glass liner (Gerstel). The following temperature program was applied during sample injection: initial temperature of 80 *^◦^*C for 30 s followed by a ramp with 12 *^◦^*C/min to 120 *^◦^*C and a second ramp with 7 *^◦^*C/min to 300 *^◦^*C and final hold for 2 min. Gas chromatographic separation was performed on a Restek Rxi-5ms column (Restek, Bad Homburg Germany). Helium was used as carrier gas with a flow rate of 1.2 mL/min. Gas chromatography was performed with the following temperature gradient: 2 min heating at 70 *^◦^*C, first temperature gradient with 5 *^◦^*C/min up to 120 *^◦^*C and hold for 30 s; subsequently, a second temperature increase of 7 *^◦^*C/min up to 350 *^◦^*C with a hold time of 2 min. The spectra were recorded in a mass range of m/z = 60 to 600 mass units with 20 spectra/s at a detector voltage of 1650 V. The GC-MS chromatograms were processed with ChromaTOF software (LECO). Mass spectra data were extracted from the Chromatof software using proprietary methods and processed using the software tool MetMax version 1.0.1.12 (MPIMP, Golm)(65) and in-house custom R scripts.

### Free nucleotide quantification

Intracellular nucleotide pools were quantified in cell lines from extracted polar metabolites using direct-infusion MS. Sample preparation was a previously described (66). Measurement of nucleotides was performed on a TSQ Quantiva triple quadrupole mass spectrometer (Thermo Fisher) coupled to a Triversa Nanomate nanoESI ion source (spray voltage: 1.5 kV, head gas pressure: 0.5 psi) and argon used as collision gas (pressure: 1.5 mTorr). FWHM Resolution for both Q1 and Q3 was set at 0.7. Data acquisition was run for 3 min per sample, using a cycle time of 3.3 sec and total acquisition of 55 SRM scans for each nucleotide in negative mode. Samples were injected in 3 technical replicates, the median of the sum of the two transitions for each of the nucleotides were calculated for analysis. The software Xcalibur (Thermo Fisher) was used to manually check the quality of each measurement, scans which presented poor signal acquisition or stalling in scanning were omitted. Data was further processed with an OpenMS package, and using in-house developed R scripts.

### Immunofluorescence imaging

Cells were seeded on circular glass slides in 6-well plates at 1e+5 cells per well. Media was exchanged for treatment with 1X PBS or 1 mM L-GA and incubated for 24 hrs. After treatment media was removed and glass slides were washed with 500 µL 1X PBS. PBS was aspirated and 500 µL of 100% ice-cold acetone was added to each well containing the slides for 10 mins. Slides were then washed 3 times with PBS. 50 µL Cytopainter Phalloidin-iFluor 488 (10X) (ab176753, Abcam) and 50 µL of Deoxyribonuclease I, Alexa Fluor 594 (1mg/ml) (D12372, Thermo Fischer) was added to each well and incubated for 20 mins in the dark. Following the staining, glass slides were washed 3 times with 1X PBS and stored at 4*^◦^*C before scanning with a Keyence bz-x700 microscope at 594nm and 488nm. Exposure times were maintained between treatment conditions. Images were processed with ImageJ 1.8.0 software. For unstained cells, cells were imaged via phase contrast microscopy at 10X magnification on a Nikon Eclipse TS2 microscope.

### Cell Cycle Analysis

Cells were seeded at 1.5e+6 cells per 10 cm plate and allowed to settle for 24 hrs. Following treatment, cells were trypsinised centrifuged for 5min at 250 x *g*, 4 *^◦^*C. The supernatant was aspirated and discarded. The cell pellet was resuspended in 1ml ice-cold 1X PBS before centrifuging for 5min at 250 x *g*, 4 *^◦^*C. The cell pellet was resuspended with 1 ml of ice-cold 70% ethanol in a drop-wise fashion while gently vortexing. Cells were fixed overnight at 4 *^◦^*C. Following fixing, cells were resuspened in 500 µL 1X PBS and 25 µL RNase A solution (10mg/ml) (Qiagen). Following incubation, 15 µL of propidium iodide (PI) (1mg/ml) (Thermo FIsher) and incubated in the dark for 15 mins at 4 *^◦^*C. PI Stained cells were transferred to fluorescence-activated cell sorting (FACS)-tubes (Sigma-Aldrich), for Fluorescence-activated cell sorting (FACS) on a BD LSR Fortessa II analyzer (Becton Dickinson) running BD FACSDIVA software v6.0 (BD Bioscience). Cell doublets were excluded by the evaluation of the propidium iodide area (PI-A) versus propidium iodide width (PI-W) density plot. Cell debris was excluded by analysing the forward scatter area and side scatter area (FSC-Area versus SSC-Area). The distinct cell cycle phases: sub-G1, G1, S and G2/M, were determined in a propidium iodide histogram. Further data analysis was performed with the FlowJo-X software (BD Bioscience, v10.0.7r2).

### Reactive oxygen species quantification

Cells were seeded at 2.5e+4 cells per well in a black-walled transparent bottom 96-well plate (Corning). Cells were allowed to attach overnight then washed once with 1X buffer provided with the DCFDA / H2DCFDA - Cellular ROS Assay Kit (ab113851). Cells were stained with 25 µL DCFDA in 1X Buffer for 45 mins at 37 *^◦^*C in an incubator. Staining media was removed and media was replaced with the drug treatment. Following 4 hrs incubation plates were scanned at Ex/Em: 485/535 nm on a Spectramax iD microplate reader (Molecular Devices). Relative fluorescence was determined relative change to control after background subtraction.

### iCelligence cell growth analysis

Cells were seeded at 1.5e+6 cells per 10 cm plate and allowed to settle for 24 hrs. Cells were optimised on the iCelligence™ system for a seeding density which provided optimal log-phase growth after 24 hrs from seeding. For both IMR-32 and SH-SY5Y a cell seeding density of 1.5e+4 per well was found to be optimal. The iCelligence™ 16-well plate was measured for background impedance and electrical contacts were checked. Cells were resuspended to a concentration of 1.5e+5 cells/ml in fresh media, 100 µL of cell suspension was added to each of the 16-wells of the iCelligence™ plate. Each of the wells were then adjusted to 200 µL with fresh media. The plate was left for 30 mins at room temperature. The plate was then inserted into the iCelligence™ device and cultivated in an incubator at 37 *^◦^*C, 5% CO_2_, 21% O_2_ and 85% relative humidity. After 24 hrs media was exchanged in each well to correspond with treatments. Impedance readings were taken in triplicate from each well every 15 mins for 120 hrs. Data was analysed with iCelligence™ DA software (ACEA Biosciences, Inc.)

### Statistics analysis

Statistical analysis were performed using the RStudio Desktop software (version 4.2.1). All data are presented as mean *±* SEM (standard error of the mean) unless specified otherwise. Student T-tests were employed for analysis between groups, normality was not tested for. To test significance between multiple groups, Anova and Tukey HSD post-hoc tests were performed. The significance levels were set at: ns *p ≥* 0.05; * *p ≤* 0.05; ** *p ≤* 0.01; *** *p ≤* 0.001; **** *p ≤* 0.0001

## Acknowledgements

We are indebted to our colleagues: J. Grobe, D. Tibutius, J. Mothes, M. Lodrini and others past and present in the Kempa and Deubzer labs. Many thanks go to C. Zasada and T. Opialla for their support with data analysis and mass spectrometry expertise. We would also like to thank A. Eggert for the continued support. Funding was provided by the TERMINATE-NB consortium (1.1.4.4, CRG-04) to S.K and H. D. Translational PhD project grant to H.D. and S. K., by German Cancer Aid funding for the ENABLE consortium [Grantno.70112951]. Bundesministerium für Bildung und Forschung funding MSTARS (Multimodal Clinical Mass Spectrometry to Target Treatment Resistance) to S.K. Helmholtz Foundation (ZT-0026) by the AMPRO consortium S.K. and Sander foundation to S.K.. Additional project funding from Charité and MDC.

## Author contributions

M.F. designed and performed experiments, analysed the data and wrote the manuscript. R.K. and L.R. performed experiments challenging L-GA with NAC. G.M. assisted in proteomic and nucleotide analysis. S.B. and B.A. assisted with pSIRM experiments. A.S. provided L-GA IC50s and M.P. provided data analysis and manuscript writing support. S.K. and H.D. supervised the work and contributed with resources, project administration and funding acquisition. S.K. conceived the project.

## References

1. B. Mendel, “Krebszelle und glycerinaldehyd,” Klinische Wochenschrift, 1929.

2. O. Warburg, K. Posener, and E. Negelein, “The metabolism of cancer cells,” Biochem Z, 1924.

3. O. Warburg, K. Gawehn, A. W. Geissler, and S. Lorenz, “Über heilung von mäuse-ascites-krebs durch d-und l-glycerinaldehyd,” Z. Klin. Chem, 1963.

4. L. H. Stickland, “The inhibition of glucolysis by glyceraldehyde,” Biochemical Journal, vol. 35, pp. 859–871, 9 1941.

5. D. M. Needham, L. Siminovitch, and S. M. Rapkine, “On the mechanism of the inhibition of glycolysis by glyceraldehyde.,” The Biochemical journal, vol. 49, pp. 113–124, 1951.

6. L. R. Bennett and F. E. Connon, “Comparative effects of dl-glyceraldehyde, 6-mercaptopurine, methotrexate and 5-fluorouracil on the ehrlich ascites carcinoma in vivo,” International Journal of Cancer, 1966.

7. J. Needham and H. Lehmann, “Intermediary carbohydrate metabolism in embryonic life,” Biochemical Journal, 1937.

8. E. Baer and H. O. Fischer, “Preparation of 1-glyceric aldehyde,” Science, vol. 88, p. 108, 7 1938.

9. K. N. Prasad, “Effect of dl-glyceraldehyde on mouse neuroblastoma cells in culture,” Cancer Research, 1972.

10. P. Schauder, C. McIntosh, J. Arends, R. Arnold, H. Frerichs, and W. Creutzfeldt, “Somatostatin and insulin release from isolated rat pancreatic islets in response to d-glucose, l-leucine, -ketoisocaproic acid or d-glyceraldehyde: Evidence for a regulatory role of adenosine-3,5-cyclic monophosphate,” Biochemical and Biophysical Research Communications, 1977.

11. S. Taniguchi, M. Okinaka, K. Tanigawa, and I. Miwa, “Difference in mechanism between glyceraldehyde- and glucose-induced insulin secretion from isolated rat pancreatic islets,” Journal of Biochemistry, 2000.

12. G. Kroemer and J. Pouyssegur, “Tumor cell metabolism: Cancer’s achilles’ heel,” Cancer Cell, vol. 13, pp. 472–482, 6 2008.

13. M. E. Irwin, N. R.-D. Valle, and J. Chandra, “Redox control of leukemia: From molecular mechanisms to therapeutic opportunities,” 4 2013.

14. Y. M. Go, J. D. Chandler, and D. P. Jones, “The cysteine proteome,” Free Radical Biology and Medicine, vol. 84, pp. 227–245, 5 2015.

15. K. K. Matthay, J. M. Maris, G. Schleiermacher, A. Nakagawara, C. L. Mackall, L. Diller, and W. A. Weiss, “Neuroblastoma,” Nature Reviews Disease Primers, 2016.

16. N. K. V. Cheung, J. Zhang, C. Lu, M. Parker, A. Bahrami, S. K. Tickoo, A. Heguy, A. S. Pappo, S. Federico, J. Dalton, I. Y. Cheung, L. Ding, R. Fulton, J. Wang, X. Chen, J. Becksfort, J. Wu, C. A. Billups, D. Ellison, E. R. Mardis, R. K. Wilson, J. R. Downing, and M. A. Dyer, “Association of age at diagnosis and genetic mutations in patients with neuroblastoma,” JAMA - Journal of the American Medical Association, 2012.

17. D. L. Baker, M. L. Schmidt, S. L. Cohn, J. M. Maris, W. B. London, A. Buxton, D. Stram, R. P. Castleberry, H. Shimada, A. Sandler, R. C. Shamberger, A. T. Look, C. P. Reynolds, R. C. Seeger, and K. K. Matthay, “Outcome after reduced chemotherapy for intermediate-risk neuroblastoma,” New England Journal of Medicine, 2010.

18. S. B. Whittle, V. Smith, E. Doherty, S. Zhao, S. McCarty, and P. E. Zage, “Overview and recent advances in the treatment of neuroblastoma,” 2017.

19. T. Wang, L. Liu, X. Chen, Y. Shen, G. Lian, N. Shah, A. M. Davidoff, J. Yang, and R. Wang, “Mycn drives glutaminolysis in neuroblastoma and confers sensitivity to an ros augmenting agent article,” Cell Death and Disease, 2018.

20. H. Takahashi, P. Oanh, T. Tran, E. Leroy, J. S. Harmon, Y. Tanaka, and R. P. Robertson, “D-glyceraldehyde causes production of intracellular peroxide in pancreatic islets, oxidative stress, and defective beta cell function via non-mitochondrial pathways,” The Journal of biological chemistry, vol. 279, p. 2020, 2004.

21. U. Raudvere, L. Kolberg, I. Kuzmin, T. Arak, P. Adler, H. Peterson, and J. Vilo, “G:profiler: A web server for functional enrichment analysis and conversions of gene lists (2019 update),” Nucleic Acids Research, vol. 47, pp. W191–W198, 7 2019.

22. S. M. Maswoswe, F. Daneshmand, and D. R. Davies, “Metabolic effects of d-glyceraldehyde in isolated hepatocytes,” Biochemical Journal, vol. 240, pp. 771–776, 1986.

23. E. Messina, P. Gazzaniga, V. Micheli, M. R. Guaglianone, S. Barbato, S. Morrone, L. Frati, A. M. Aglianó, and A. Giacomello, “Guanine nucleotide depletion triggers cell cycle arrest and apoptosis in human neuroblastoma cell lines,” International Journal of Cancer, 2004.

24. M. Camici, M. Garcia-Gil, R. Pesi, S. Allegrini, and M. G. Tozzi, “Purine-metabolising enzymes and apoptosis in cancer,” Cancers, 2019.

25. D. E. Atkinson and G. M. Walton, “Adenosine triphosphate conservation in metabolic regulation. rat liver citrate cleavage enzyme.,” Journal of Biological Chemistry, 1967.

26. S. Mandal, W. A. Freije, P. Guptan, and U. Banerjee, “Metabolic control of g1-s transition: Cyclin e degradation by p53-induced activation of the ubiquitin-proteasome system,” Journal of Cell Biology, 2010.

27. M. R. Buchakjian and S. Kornbluth, “The engine driving the ship: Metabolic steering of cell proliferation and death,” Nature Reviews Molecular Cell Biology, vol. 11, pp. 715–727, 2010.

28. J. Kalucka, R. Missiaen, M. Georgiadou, S. Schoors, C. Lange, K. D. Bock, M. Dewerchin, and P. Carmeliet, “Metabolic control of the cell cycle,” Cell Cycle, vol. 14, pp. 3379–3388, 2015.

29. M. Salazar-Roa and M. Malumbres, “Fueling the cell division cycle,” Trends in Cell Biology, 2017.

30. M. Pietzke, “Analysis of the metabolic control of cell growth using stable isotope resolved metabolomics,” 2015.

31. M. Pietzke, C. Zasada, S. Mudrich, and S. Kempa, “Decoding the dynamics of cellular metabolism and the action of 3-bromopyruvate and 2-deoxyglucose using pulsed stable isotope-resolved metabolomics,” Cancer Metabolism, vol. 2, p. 9, 2014.

32. P. V. Hornbeck, B. Zhang, B. Murray, J. M. Kornhauser, V. Latham, and E. Skrzypek, “Phosphositeplus, 2014: mutations, ptms and recalibrations,” Nucleic acids research, vol. 43, pp. D512–D520, 1 2015.

33. C. J. Martyniuk, B. Fang, J. M. Koomen, T. Gavin, L. Zhang, D. S. Barber, and R. M. Lopachin, “Molecular mechanism of glyceraldehyde-3-phosphate (gapdh) inactivation by,-unsaturated carbonyl derivatives.,” Chemical research in toxicology, vol. 24, p. 2302, 12 2011.

34. M. J. MacDonald, F. W. Chaplen, C. K. Triplett, Q. Gong, and H. Drought, “Stimulation of insulin release by glyceraldehyde may not be similar to glucose,” Archives of Biochemistry and Biophysics, 2006.

35. J. V. D. Reest, S. Lilla, L. Zheng, S. Zanivan, and E. Gottlieb, “Proteome-wide analysis of cysteine oxidation reveals metabolic sensitivity to redox stress,” Nature Communications 2018 9:1, vol. 9, pp. 1–16, 4 2018.

36. H. A. Lardy, V. D. Wiebelhaus, and K. M. Mann, “The mechanism by which glyceraldehyde inhibits glycolysis.,” The Journal of biological chemistry, 1950.

37. J. Phan, E. Mahdavian, M. C. Nivens, W. Minor, S. Berger, H. T. Spencer, R. B. Dunlap, and L. Lebioda, “Catalytic cysteine of thymidylate synthase is activated upon substrate binding†,‡,” Biochemistry, vol. 39, pp. 6969–6978, 6 2000.

38. P. Nordlund and P. Reichard, “Ribonucleotide reductases,” annurev.biochem., vol. 75, pp. 681–706, 6 2006.

39. H. Alborzinia, A. F. Flórez, S. Kreth, L. M. Brückner, U. Yildiz, M. Gartlgruber, D. I. Odoni, G. Poschet, K. Garbowicz, C. Shao, C. Klein, J. Meier, P. Zeisberger, M. Nadler-Holly, M. Ziehm, F. Paul, J. Burhenne, E. Bell, M. Shaikhkarami, R. Würth, S. A. Stainczyk, E. M. Wecht, J. Kreth, M. Büttner, N. Ishaque, M. Schlesner, B. Nicke, C. Stresemann, M. Llamazares-Prada, J. H. Reiling, M. Fischer, I. Amit, M. Selbach, C. Herrmann, S. Wölfl, K. O. Henrich, T. Höfer, A. Trumpp, and F. Westermann, “Mycn mediates cysteine addiction and sensitizes neuroblastoma to ferroptosis,” Nature Cancer 2022 3:4, vol. 3, pp. 471–485, 4 2022.

40. I. A. Manke, A. Nguyen, D. Lim, M. Q. Stewart, A. E. Elia, and M. B. Yaffe, “Mapkap kinase-2 is a cell cycle checkpoint kinase that regulates the g2/m transition and s phase progression in response to uv irradiation,” Molecular cell, vol. 17, pp. 37–48, 1 2005.

41. S. Macip, A. Kosoy, S. W. Lee, M. J. O’Connell, and S. A. Aaronson, “Oxidative stress induces a prolonged but reversible arrest in p53-null cancer cells, involving a chk1-dependent g2 checkpoint,” Oncogene 2006 25:45, vol. 25, pp. 6037–6047, 5 2006.

42. A. Krystyniak, C. Garcia-Echeverria, C. Prigent, and S. Ferrari, “Inhibition of aurora a in response to dna damage,” Oncogene, vol. 25, pp. 338–348, 1 2006.

43. S. G. DuBois, J. D. Kremer, B. D. Wilde, C. Jacobsen, I. Aerts, Y. P. Mosse, J. M. Maris, A. Lithio, A. Gosberg, C. Banks, C. Tate, M. Dowless, X. Gong, L. Stancato, K. M. Bell-McGuinn, J. R. Park, A. D. Pearson, and A. Marachelian, “A phase i study of aurora kinase a inhibitor ly3295668 erbumine as a single agent and in combination in patients with relapsed/refractory neuroblastoma.,” vol. 38, 5 2020.

44. H. Pelicano, D. Carney, and P. Huang, “Ros stress in cancer cells and therapeutic implications,” Drug Resistance Updates, vol. 7, pp. 97–110, 4 2004.

45. H. Mizutani, S. Tada-Oikawa, Y. Hiraku, M. Kojima, and S. Kawanishi, “Mechanism of apoptosis induced by doxorubicin through the generation of hydrogen peroxide,” Life Sciences, vol. 76, pp. 1439–1453, 2 2005.

46. R. Marullo, E. Werner, N. Degtyareva, B. Moore, G. Altavilla, S. S. Ramalingam, and P. W. Doetsch, “Cisplatin induces a mitochondrial-ros response that contributes to cytotoxicity depending on mitochondrial redox status and bioenergetic functions,” PLoS ONE, 2013.

47. U. Ozer, K. W. Barbour, S. A. Clinton, and F. G. Berger, “Oxidative stress and response to thymidylate synthase-targeted antimetabolites,” Molecular Pharmacology, vol. 88, pp. 970–981, 12 2015.

48. M. Uetaki, S. Tabata, F. Nakasuka, T. Soga, and M. Tomita, “Metabolomic alterations in human cancer cells by vitamin c-induced oxidative stress,” Scientific reports, vol. 5, 9 2015.

49. L. S. Ihrlund, E. Hernlund, O. Khan, and M. C. Shoshan, “3-bromopyruvate as inhibitor of tumour cell energy metabolism and chemopotentiator of platinum drugs,” Molecular Oncology, 2008.

50. Y. H. Ko, H. A. Verhoeven, M. J. Lee, D. J. Corbin, T. J. Vogl, and P. L. Pedersen, “A translational study “case report” on the small molecule “energy blocker” 3-bromopyruvate (3bp) as a potent anticancer agent: from bench side to bedside,” Journal of Bioenergetics and Biomembranes 2012 44:1, vol. 44, pp. 163–170, 2 2012.

51. L. E. Raez, K. Papadopoulos, A. D. Ricart, E. G. Chiorean, R. S. Dipaola, M. N. Stein, C. M. R. Lima, J. J. Schlesselman, K. Tolba, V. K. Langmuir, S. Kroll, D. T. Jung, M. Kurtoglu, J. Rosenblatt, and T. J. Lampidis, “A phase i dose-escalation trial of 2-deoxy-d-glucose alone or combined with docetaxel in patients with advanced solid tumors,” Cancer chemotherapy and pharmacology, vol. 71, pp. 523–530, 2 2013.

52. R. Kapoor, D. B. Gundpatil, B. L. Somani, T. K. Saha, S. Bandyopadhyay, and P. Misra, “Anticancer effect of dl-glyceraldehyde and 2-deoxyglucose in ehrlich ascites carcinoma bearing mice and their effect on liver, kidney and haematological parameters,” Indian Journal of Clinical Biochemistry, 2014.

53. S. Ganapathy-Kanniappan, M. Vali, R. Kunjithapatham, M. Buijs, L. Syed, P. Rao, S. Ota, B. Kwak, R. Loffroy, and J. Geschwind, “3-bromopyruvate: A new targeted antiglycolytic agent and a promise for cancer therapy,” Current Pharmaceutical Biotechnology, vol. 11, pp. 510–517, 6 2010.

54. L. MacChioni, M. Davidescu, M. Sciaccaluga, C. Marchetti, G. Migliorati, S. Coaccioli, R. Roberti, L. Corazzi, and E. Castigli, “Mitochondrial dysfunction and effect of antiglycolytic bromopyruvic acid in gl15 glioblastoma cells,” Journal of bioenergetics and biomembranes, vol. 43, pp. 507–518, 10 2011.

55. E. G. Konstantakou, G. E. Voutsinas, A. D. Velentzas, A. S. Basogianni, E. Paronis, E. Balafas, N. Kostomitsopoulos, K. N. Syrigos, E. Anastasiadou, and D. J. Stravopodis, “3-brpa eliminates human bladder cancer cells with highly oncogenic signatures via engagement of specific death programs and perturbation of multiple signaling and metabolic determinants,” Molecular cancer, vol. 14, 7 2015.

56. M. C. Coleman, C. R. Asbury, D. Daniels, J. Du, N. Aykin-Burns, B. J. Smith, L. Li, D. R. Spitz, and J. J. Cullen, “2-deoxy-d-glucose causes cytotoxicity, oxidative stress, and radiosensitization in pancreatic cancer,” Free Radical Biology and Medicine, 2008.

57. P. Muley, A. Olinger, and H. Tummala, “2-deoxyglucose induces cell cycle arrest and apoptosisin colorectal cancer cells independent of its glycolysis inhibition,” Nutrition and Cancer, 2015.

58. N. Brock and T. Niekamp, “Zur frage der cytostatischen wirksamkeit von d-glycerinaldehyd,” Zeitschrift für Krebsforschung, 1965.

59. A. D. Pearson, C. R. Pinkerton, I. J. Lewis, J. Imeson, C. Ellershaw, and D. Machin, “High-dose rapid and standard induction chemotherapy for patients aged over 1 year with stage 4 neuroblastoma: a randomised trial,” The Lancet Oncology, 2008.

60. J. R. Yates, “The revolution and evolution of shotgun proteomics for large-scale proteome analysis,” Journal of the American Chemical Society, 2013.

61. S. Tyanova, T. Temu, P. Sinitcyn, A. Carlson, M. Y. Hein, T. Geiger, M. Mann, and J. Cox, “The perseus computational platform for comprehensive analysis of (prote)omics data,” Nature Methods, 2016.

62. R. Ihaka and R. Gentleman, “R: A language for data analysis and graphics,” Journal of Computational and Graphical Statistics, vol. 5, pp. 299–314, 1996.

63. Y. Perez-Riverol, J. Bai, C. Bandla, D. García-Seisdedos, S. Hewapathirana, S. Kamatchinathan, D. J. Kundu, A. Prakash, A. Frericks-Zipper, M. Eisenacher, M. Walzer, S. Wang, A. Brazma, and J. A. Vizcaíno, “The pride database resources in 2022: a hub for mass spectrometry-based proteomics evidences,” Nucleic Acids Research, vol. 50, p. D543, 1 2022.

64. C. Zasada and S. Kempa, Quantitative analysis of cancer metabolism: From pSIRM to MFA, pp. 207–220. Springer, 2016.

65. S. Kempa, J. Hummel, T. Schwemmer, M. Pietzke, N. Strehmel, S. Wienkoop, J. Kopka, and W. Weckwerth, “An automated gcxgc-tof-ms protocol for batch-wise extraction and alignment of mass isotopomer matrixes from differential13c-labelling experiments: A case study for photoautotrophic-mixotrophic grown chlamydomonas reinhardtii cells,” Journal of Basic Microbiology, 2009.

66. P. Lorkiewicz, R. M. Higashi, A. N. Lane, and T. W.-M. Fan, “High information throughput analysis of nucleotides and their isotopically enriched isotopologues by direct-infusion fticr-ms,” Metabolomics, vol. 8, pp. 930–939, 10 2012.

